# A human missense variant in BSX decouples circadian behaviour from metabolic rhythmicity while preserving lifespan in mice

**DOI:** 10.64898/2025.12.05.692520

**Authors:** Anna-Corina Treier, Christian Klasen, Madlen Kramer, Deborah Schmidt, Franziska Block, Janine Gebhardt, Fabio Rojas Rusak, I. Sadaf Farooqi, Mathias Treier

## Abstract

Mammalian circadian organisation is orchestrated by a light-entrained master pacemaker in the suprachiasmatic nucleus (SCN) and SCN-independent, food- and metabolism-responsive oscillators in the brain and periphery; feeding can entrain these systems, enabling anticipation of feeding time. Here, we show that a human missense variant (Q159K) in the homeodomain of the brain-specific transcription factor BSX causes profound loss of circadian behavioural rhythms and food anticipatory activity (FAA) in mice. Whole-brain imaging reveals failure of the compact dorsomedial hypothalamus (DMHc) to form and the absence of histaminergic neurons in the tuberomammillary nucleus (TMN) in *Bsx^Q159K/Q159K^* mice. Consequently, under light-dark (LD) and constant darkness (DD), sleep-wake cycles and circadian locomotor rhythmicity are profoundly lost, whereas total wake time and spontaneous locomotor activity are preserved, with daily caloric intake substantially shifted from the dark to the light cycle. Therefore, in DD, metabolic recordings reveal a rhythmic respiratory exchange ratio (RER) pattern that differs substantially from that of wild-type mice and is markedly attenuated under LD conditions. In addition, *Bsx^Q159K/Q159K^* mice lack FAA during restricted feeding and fail to increase locomotor activity upon food deprivation. Despite the loss of circadian behavioural rhythms and altered daily feeding behaviour, *Bsx^Q159K/Q159K^*mice maintain metabolic homeostasis, a lean body composition into advanced age, and a normal lifespan under ad libitum feeding and controlled LD. These findings provide genetic evidence that circadian organization is not universally required for metabolic homeostasis, suggesting instead that circadian rhythmicity is an adaptive feature advantageous primarily under diurnal conditions.

## Introduction

Animals rely on circadian clocks, endogenous oscillators that coordinate 24 h physiological and behavioural processes, including sleep-wake cycles, locomotion, and feeding [1–5]. In mammals, a light-entrained pacemaker in the suprachiasmatic nucleus (SCN) aligns internal time with the external light-dark cycle via retinal input [1, 6]. Restricted feeding can entrain clocks outside the SCN, thereby uncoupling circadian oscillators in peripheral tissues from the SCN, rephasing peripheral oscillators and generating food-anticipatory activity (FAA) [7–9]. Whether FAA arises from a single food-entrainable oscillator (FEO) or a distributed network of multiple interacting oscillators remains debated [10, 11].

The dorsomedial hypothalamus (DMH) receives SCN output via the subparaventricular zone and is implicated in the organisation of sleep-wake and feeding rhythms [12]. Under restricted feeding, *Per1/Per2* (*period1/period2*) rhythms and *Bmal1* (*basic helix-loop-helix ARNT like 1*) expression have been reported in the compact DMH (DMHc), a densely packed subregion with strong connections to the preoptic area (POA), paraventricular (PVN) and lateral hypothalamic (LH) centres [13, 14]. Molecular markers enriched in DMHc include *Ppp1r17* (protein phosphatase 1 regulatory subunit 17), *Grp* (*gastrin-releasing peptide*), *Cck* (*cholecystokinin*), *Pdyn* (*Prodynorphin*) and *Bsx* itself. DMHc neurons project to the PVN where they have been shown, upon activation, to suppress food intake [15]. Furthermore, DMHc PPP1R17-positive neurons have also been implicated in limiting food intake and body weight of *ob* (leptin-deficient) mice and, by knockdown of *Ppp1r17* itself in the DMHc, in influencing WAT function and lifespan [16, 17].

Histaminergic neurons of the tuberomammillary nucleus (TMN) promote wakefulness and locomotor activation via widespread forebrain projections [18, 19]. Together, DMHc and TMN constitute critical nodes coupling circadian timing to arousal and metabolism [5, 20]. However, the necessity of DMH (and DMHc) for FAA remains controversial, with lesion studies yielding contradictory results [21–24]. The brain-specific homeobox factor BSX, with a Q50-class homeodomain, contributes to hypothalamic developmental programmes and is required in the arcuate nucleus (ARC) for normal AgRP/NPY expression [25]. The ARC itself is central to the regulation of body weight, feeding behavior, growth, and reproduction, and its neuronal specification establishes long-term metabolic balance [26].

We hypothesised that altering BSX protein activity would remodel hypothalamic circuitry controlling behavioural and metabolic rhythms. Here we show that a very rare human missense variant which alters the homeodomain residue at position 50 in BSX, from glutamine (Q50) to lysine (K50) (Q159K), induces profound anatomical reorganisation of the hypothalamus, including loss of the DMHc and of histidine decarboxylase (HDC) positive TMN neurons, eliminating key components that link circadian timing to arousal and metabolism. As a consequence, *Bsx^Q159K/Q159K^* mice exhibit profound loss of circadian behavioural rhythms (sleep-wake, locomotion and feeding) under LD and DD, yet retain a detectable ∼24 h RER rhythm in DD that is attenuated in LD, and maintain metabolic homeostasis, leanness and normal lifespan under ad libitum conditions. Additionally, they show diminished food-seeking behavior during food deprivation and absence of FAA. These findings provide genetic evidence that DMHc and histaminergic neurons are critical nodes for behavioural circadian output and reveal a dissociation between behavioural and metabolic timing, indicating that temporal organisation can be conditionally dispensable for metabolic homeostasis in stable environments.

## Results

### Generation of mouse strains carrying human BSX amino acid missense variants

To investigate the possible roles of BSX in mammalian physiology and behaviour, we took advantage of the fact that human and mouse BSX proteins are nearly identical, with their homeodomain protein sequences matching 100%.

We selected the following human BSX amino acid (AA) missense variants from gnomAD v4.1 [27]: Proline11Alanine, Glycine55Arginine, Threonine145Lysine, Glutamine173Proline, Glutamate183Glycine, Histidine231Glutamine. In addition, we searched for rare missense variants in *BSX* in 927 exomes from children with severe early-onset obesity (defined as Body Mass Index (BMI) standard deviation score > 3, onset before age 10 years) and of White British ancestry, recruited to the Genetics of Obesity Study (GOOS) cohort [28]. We identified one heterozygous rare variant (Q159K) in the highly constrained DNA-binding domain that was absent from almost 800,000 exomes/genomes in gnomAD v4.1. The missense variant was identified in a 15-year-old girl with severe obesity (BMI SD score 3.0) and learning difficulties, and arose *de novo* (i.e., absent in the DNA from both parents). This missense variant changes glutamine at position 50 of the BSX homeodomain, characteristic of the Q50-type homeodomain family, into lysine found in the K50-type homeodomain family, which determines DNA-binding specificity [29–31].

Using CRISPR/Cas9 technology, we introduced each AA missense variant on its own into C57BL/6J mice to create independent mouse strains carrying the respective AA missense variant. We backcrossed all mouse strains for one generation and confirmed, by sequencing, the successful introduction of the AA missense variant and the absence of any other alterations in *Bsx*. Subsequently, for each human AA missense variant, homozygous carrier mouse strains were generated. Six of seven strains did not show obvious body size and weight differences compared to their C57BL/6J wild-type littermates.

Whereas *Bsx^Q159K/WT^* mice were indistinguishable from their wild-type littermates, *Bsx^Q159K/Q159K^*male and female mice homozygous for the human Glutamine159Lysine (Q159K) variant exhibited, unexpectedly and in contrast to the human female heterozygous carrier, approximately 25% lower body weight at three months of age, due to reduced body length and lean mass compared with their wild-type littermates (Suppl. Fig. 1A,B and data not shown). Despite smaller body size, fertility was not affected in *Bsx^Q159K/Q159K^* mice, as breeding of *Bsx^Q159K/Q159K^* males with *Bsx^Q159K/Q159K^* females resulted in viable offspring.

The obvious phenotypic difference between *Bsx^Q159K/Q159K^*mice and *Bsx^-/-^* knockout mice, which do not show a body size difference compared to their C57BL6/J wild-type littermates, suggested that the human Q159K missense variant does not affect mouse BSX protein expression. This was confirmed by immunohistochemistry with a BSX-specific antibody, which detected BSX^Q159K^ protein expression in the arcuate nucleus and dorsomedial hypothalamus (DMH) of *Bsx^Q159K/Q159K^* mice (Suppl. Fig. 1C), where BSX has been shown to directly regulate the orexigenic peptide-encoding *Npy* and *Agrp* genes [25]. We therefore further analysed the mouse strain with the human BSX^Q159K^ missense variant.

### Comparative light-sheet image analysis of lineage-traced and activated BSX neurons in BSX^WT^ and BSX^Q159K^ brains

BSX is specifically expressed in the brain, most prominently in the hypothalamus. Therefore, to evaluate whether changes in cell number, anatomical localisation, and/or function of BSX-expressing neurons are present in Bsx^Q159K/Q159K^ compared to Bsx^WT/WT^ brains, we employed an unbiased whole-brain imaging approach. To trace and activate BSX^WT^- or BSX^Q159K^-expressing cells throughout the respective brains, the following genetic setups were established. First, a *Bsx^Cre^* knock-in allele was generated, placing Cre recombinase at the ATG start site of the *Bsx* gene, which at the same time resulted in a *Bsx* null allele. Combining one *Bsx^Cre^*allele with either one *Bsx^WT^* or *Bsx^Q159K^* allele together with conditional alleles for Rosa-Ai14-tdTomato and R26-hM3Dq/mCitrine (the latter allowing neuronal activation by clozapine-N-oxide, CNO) [32] resulted in littermate mice with the following genotypes: *Bsx^Cre/WT^/Ai14-tdTomato/R26-hM3Dq* or *Bsx^Cre/Q159K^/Ai14-tdTomato/R26-hM3Dq*. Expression of tdTomato and Citrine in *Bsx^Cre/WT^/Ai14-tdTomato/R26-hM3Dq* mice faithfully recapitulated endogenous BSX expression as demonstrated by BSX/tdTomato and BSX/mCitrine double immunohistochemistry (data not shown). These genotypes therefore allowed us to lineage-trace BSX^WT^ or BSX^Q159K^ neurons and, at the same time, to map c-Fos distribution upon BSX^WT^ or BSX^Q159K^ neuronal network activation by CNO.

Two hours after CNO application, adult brains of the respective genotypes were paraformaldehyde-fixed, followed by tissue-clearing and 3D immunostaining with c-FOS/NeuN antibodies, and further processed for light-sheet imaging. Whole-brain light-sheet imaging was performed with direct detection of tdTomato at 561 nm, c-FOS at 647 nm, and NeuN at 488 nm. NeuN was used to facilitate the alignment of the imaged brains with the anatomical Allen Brain Atlas [33]. Whole-brain image analysis allowed us to zoom through BSX^WT^ and BSX^Q159K^ brains from all angles, (coronal, sagittal and transverse) to observe the respective BSX^tdTomato^- and/or c-FOS-positive cells in an unbiased manner (Fig. 1, videos). Overlay of whole-brain image data sets from *Bsx^Cre/WT^/Ai14-tdTomato/R26-hM3Dq* and *Bsx^Cre/Q159K^/Ai14-tdTomato/R26-hM3Dq* brains directly visualised differences in the localisation of BSX- and/or c-FOS-expressing neurons throughout BSX^WT^ versus BSX^Q159K^ brains. Zooming in on the hypothalamic region, which harbours most BSX-expressing neurons, we could readily identify regions that showed alterations in the amount and distribution of BSX- and/or c-FOS-expressing neurons. Most strikingly, the compaction of BSX^tdTomato^-positive neurons in the dorsomedial hypothalamus (DMH) was absent in BSX^Q159K^ brains. Furthermore, far fewer BSX^tdTomato^-positive neurons were seen in the ventral region of the tuberomammillary nucleus (TMN) in BSX^Q159K^ compared to BSX^WT^ brains. In contrast, c-FOS-expressing BSX positive neurons were more abundant in BSX^Q159K^ brains upon CNO activation, most prominently in the DMH and posterior hypothalamic area (PH), suggesting that part of the BSX neurons have become re-localised within the hypothalamus. To independently confirm these findings, we employed a previously generated *Bsx^H2BEGFP^* allele, which at the same time is a *Bsx* null allele, to generate the following genotypes: *Bsx^WT/H2BEGFP^* and *Bsx^Q159K/H2BEGFP^*. Imaging of rostral-caudal serial coronal sections spanning the whole DMH-TMN region again revealed that Bsx^Q159K/H2BEGFP^ lineage-traced neurons failed to compact within the DMH and were absent in the ventral TMN region (Suppl. Fig. 2). Finally, it is important to note that neither tracing study revealed any expression of *Bsx* in or close to the SCN (data not shown).

**Fig. 1:**
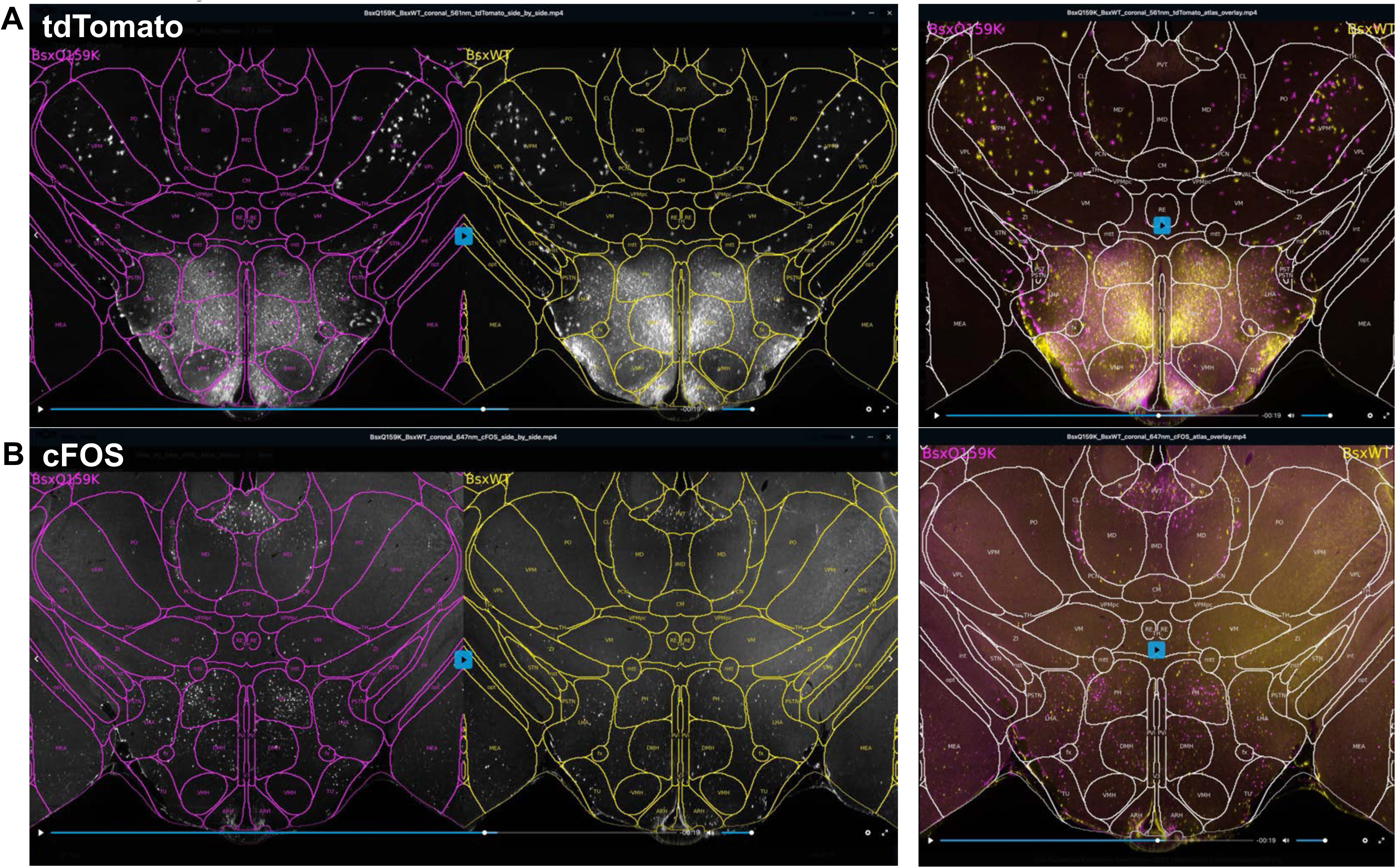
Snapshots from whole brain image video analysis of Bsx^Cre/WT^/Ai14-tdTomato/R26-hM3Dq or Bsx^Cre/Q159K^/Ai14-tdTomato/R26-hM3Dq mice. **A)** Coronal section snapshot at the level of the dorsomedial hypothalamus (DMH) of tdTomato whole brain image video analysis from Bsx^Q159K^ and Bsx^WT^ brains sidebyside (left) or overlayed (right) demonstrating loss of cellular compaction of BSX lineage traced neurons in the DMH of Bsx^Q159K^ brains. **B)** Coronal section snapshot at the level of the DMH of cFOS whole brain image video analysis from Bsx^Q159K^ and Bsx^WT^ brains sidebyside (left) or overlayed (right) demonstrating ectopic cFOS expression in BSX lineage traced neurons upon CNO application in the DMH of Bsx^Q159K^ brains.

### Alterations of hypothalamic gene expression in homozygous Bsx^Q159K^ mice

To confirm the observed anatomical differences in the DMH of BSX^Q159K/Q159K^ brains at the molecular level, we took advantage of the Allan Single-Cell Mouse Brain Atlas [34] to select the following DMH-specific marker genes for *in situ* hybrydisation (ISH): *Grp* (*gastrin-releasing peptide*), *Cck* (*cholecystokinin*), *Ppp1r17* (*protein phosphatase 1 regulatory subunit 17*), and *Bsx* itself. In wild-type mouse brains these markers give rise to a “butterfly-shaped” expression pattern representing the compact region of the DMH (DMHc) in ISH (Fig.2). *Trh* (*thyrotropin-releasing hormone*) was included as a reference marker for the region, being intermingled with, but not showing, a “butterfly-shaped” expression pattern. ISH with these marker genes, including a *tdTomato* in situ probe, recapitulated the expected “butterfly-shaped” expression pattern and confirmed that *Bsx* itself is a marker gene for the compact region of the DMH throughout the hypothalamus. In contrast, in BSX^Q159K/Q159K^ brains the “butterfly-shaped” expression patterns of the three DMHc marker genes, in particular *Ppp1r17* and *Grp*, were absent from rostral to caudal throughout the DMH of the hypothalamus (Fig.2). Together with the loss of compaction seen for the BSX^tdTomato^- or BSX^H2BEGFP^-positive cells, these data demonstrate that the compact region of the DMH, with its neuronal marker gene expression, is absent in *Bsx^Q159K/Q159K^* mice at the molecular level. With respect to the observed reduction of BSX lineage-traced tdTomato- or H2BEGFP-positive cells in the TMN, expression of *Hdc* (*histamine decarboxylase*), the signature gene within the dorsal and ventral TMN, was examined by *in situ* hybridization [35]. To our surprise, *Hdc* expression was absent throughout the TMN. To distinguish whether only *Hdc* expression is gone in the TMN or whether HDC-expressing neurons themselves are not present, we used *Prph* (*peripherin*, an intermediate filament) gene expression as a decisive molecular marker. *Prph* is expressed only in the TMN within the central nervous system and prior to *Hdc* during development, with both markers being exclusively co-expressed in the adult brain. The absence of *Prph* expression, as well as of expression of two other HDC neuronal marker genes, *Ox2r* (*orexin 2 receptor*) and Wif1 (*Wnt inhibitory factor 1*), throughout the TMN of *Bsx^Q159K/Q159K^* mice strongly suggested that not only HDC expression is lost but that *Hdc*-expressing neurons themselves are not present in BSX^Q159K/Q159K^ brains (Fig. 3).

**Fig. 2:**
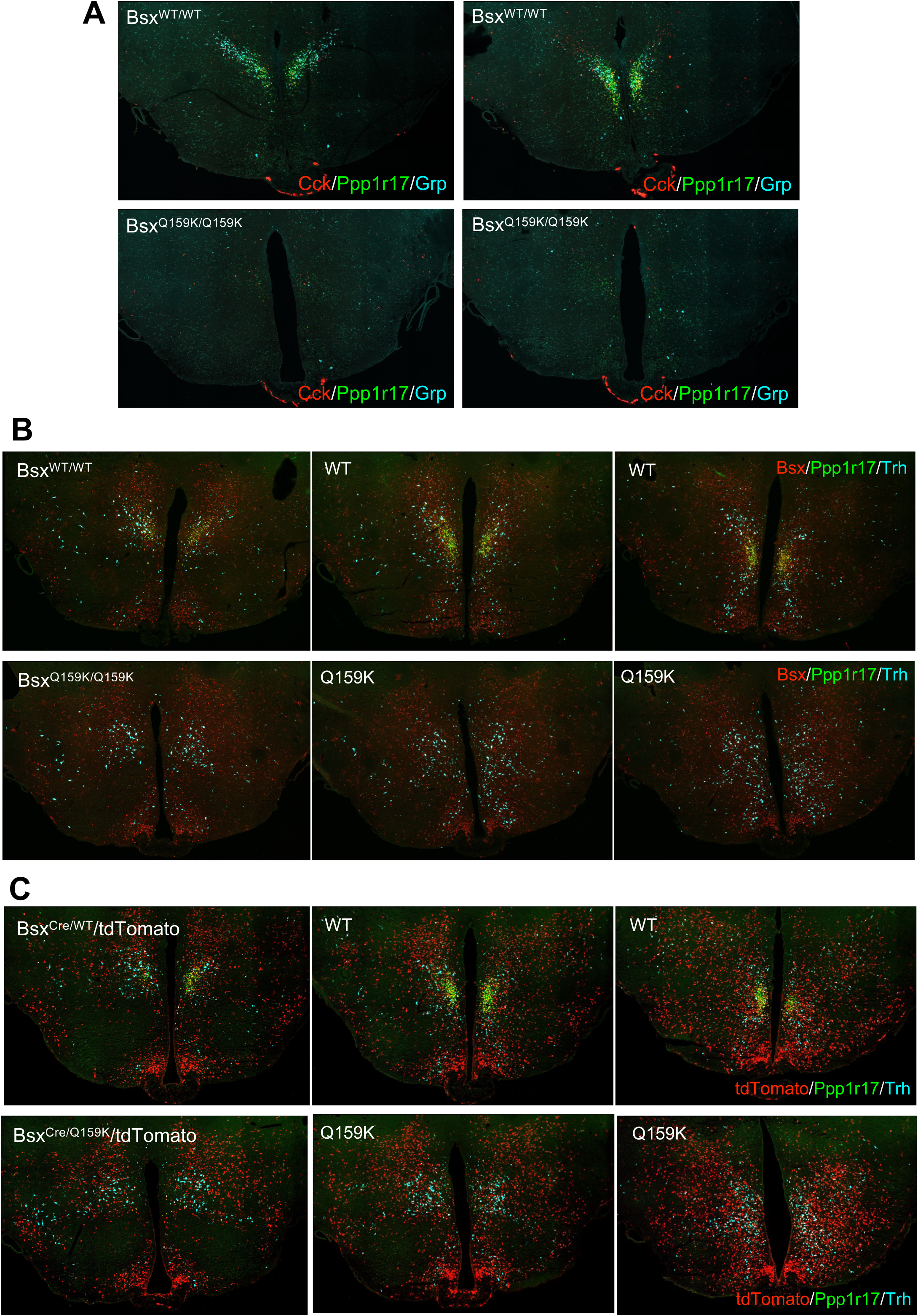
The compact region of the DMH (DMHc) is absent in Bsx^Q159K/Q159k^ mice. **A)** Representative tile scans of coronal sections of the hypothalamus at the level of the 3^rd^ ventrical from rostral (left) to caudal (right) after *in situ* hybridization (ISH) with Cck, Ppp1r17 and Grp probes from brains of BSX^WT/WT^ mice (upper sections) and Bsx^Q159K/Q159K^ mice (lower sections). Whereas all three marker genes Cck, Ppp1r17 and Grp of the compact region of the DMH (DMHc) with their characteristic “butterfly shape” expression pattern are present in BSX^WT/WT^ brains their expression is absent in Bsx^Q159K/Q159K^ brains. **B)** Representative tile scans of coronal sections of the hypothalamus at the level of the 3^rd^ ventrical from rostral (left) to caudal (right) after *in situ* hybridization (ISH) with Bsx, Ppp1r17 and Trh probes from brains of BSX^WT/WT^ mice (upper sections) and Bsx^Q159K/Q159K^ mice (lower sections). All three genes Bsx, Ppp1r17 and Trh are present in BSX^WT/WT^ brains with co-expression of Bsx and Ppp1r17 in DMHc neurons. In contrast Ppp1r17 expression, as DMHc marker gene, is absent and the optical dense expression of Bsx is gone due to the absence of the DMHc in Bsx^Q159K/Q159K^ brains, with Trh expression not majorly different between the two genotypes. **C)** Representative tile scans of coronal sections of the hypothalamus at the level of the 3^rd^ ventrical from rostral (left) to caudal (right) after *in situ* hybridization (ISH) with tdTomato, Ppp1r17 and Trh probes from brains of BSX^Cre/WT^/tdTomato mice (upper sections) and Bsx^Cre/Q159K^/tdTomato mice (lower sections) mice . All three genes tdTomato, Ppp1r17 and Trh are present in BSX^Cre/WT^/tdTomato brains with co-expression of tdTomato and Ppp1r17 in DMHc neurons. In contrast Ppp1r17 expression, as DMHc marker gene, is absent and the optical dense expression of tdTomato is gone similar to what is observed in whole brain light-sheet imaging due to the absence of the DMHc in BSX^Cre/WT^/tdTomato brains with Trh expression not majorly different between the two genotypes.

**Fig. 3:**
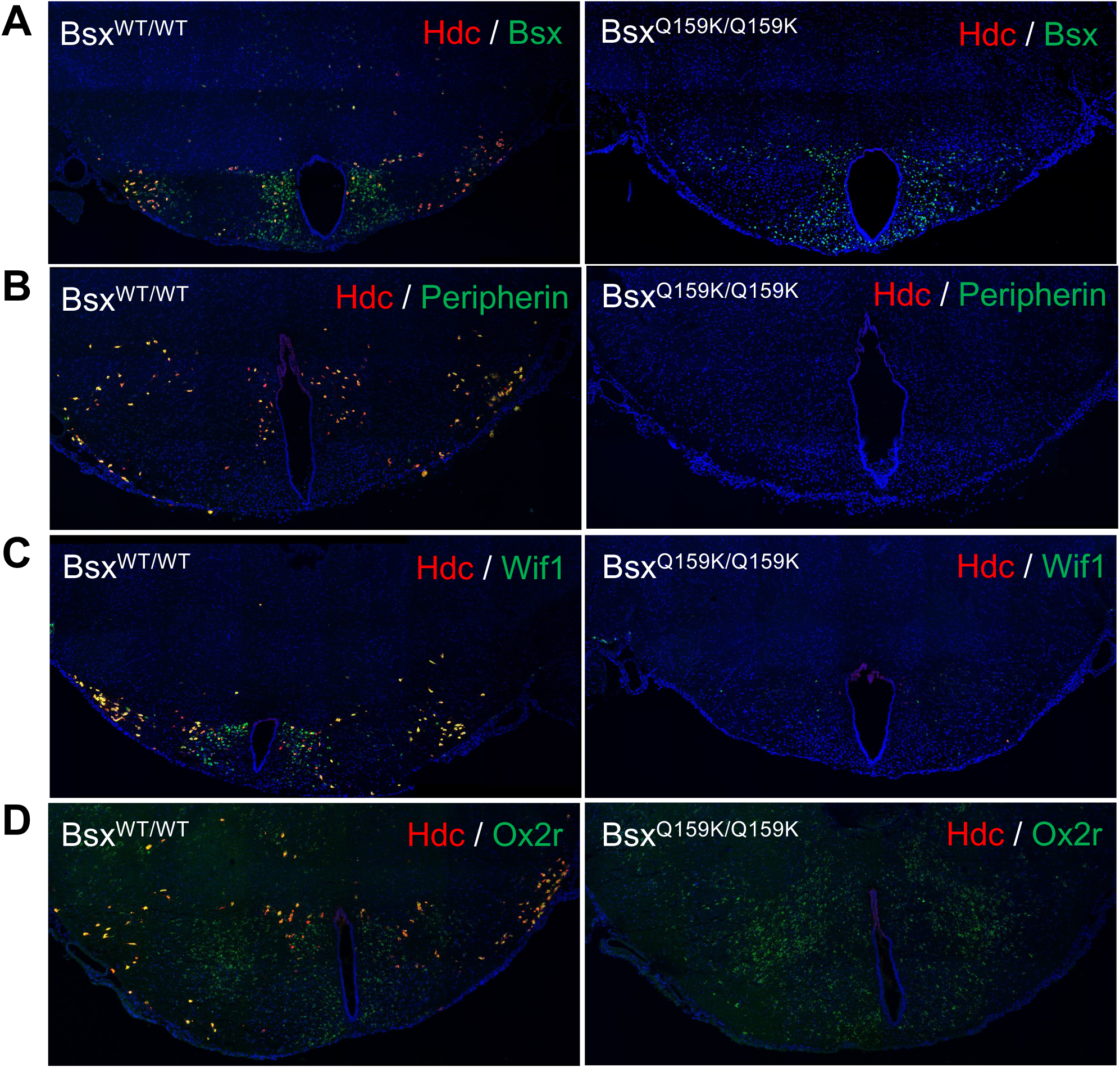
Histidine decarboxylase (HDC) positive neurons are absent in Bsx^Q159K/Q159k^ mice. **A)** Representative tile scans of coronal sections of the tuberomammillary nucleus (TMN) of the posterior hypothalamus after *in situ* hybridization (ISH) with Hdc and Bsx probes from brains of BSX^WT/WT^ mice (left) and Bsx^Q159K/Q159K^ mice (right). Hdc and Bsx are co-expressed in vTMN of BSX^WT/WT^ brains (yellow) with co-expression of both markers absent in Bsx^Q159K/Q159K^ brains consistent with the observed loss of BSX lineage traced neurons in the vTMN. **B)** Representative tile scans of coronal sections of TMN after *in situ* hybridization (ISH) with Hdc and Peripherin probes from brains of BSX^WT/WT^ mice (left) and Bsx^Q159K/Q159K^ mice (right). Hdc and Peripherin are co-expressed throughout the TMN of BSX^WT/WT^ brains (yellow) but expression of both markers is absent in Bsx^Q159K/Q159K^ brains. **C)** Representative tile scans of coronal sections of TMN after *in situ* hybridization (ISH) with Hdc and Wif1 probes from brains of BSX^WT/WT^ mice (left) and Bsx^Q159K/Q159K^ mice (right). Hdc and Wif1 are co-expressed throughout the TMN of BSX^WT/WT^ brains (yellow) but expression of both markers is absent in Bsx^Q159K/Q159K^ brains. **D)** Representative tile scans of coronal sections of TMN after *in situ* hybridization (ISH) with Hdc and Ox2r probes from brains of BSX^WT/WT^ mice (left) and Bsx^Q159K/Q159K^ mice (right). Hdc and Ox2r are co-expressed in the TMN of BSX^WT/WT^ brains (yellow) but no HDC expression is observed in Bsx^Q159K/Q159K^ brains.

The composition of cell types in the arcuate nucleus has been shown to be determined by a homeobox code, with BSX being part of it [36, 37]. Because BSX^Q159K^ protein is readily detectable in the arcuate nucleus of *Bsx^Q159K/Q159K^* mice, we wondered whether the AA substitution would affect the cell-type composition of the arcuate nucleus. We have previously shown that BSX is important for expression of the two orexigenic peptide-encoding genes *Npy* and *Agrp*, as well as the responsiveness of AgRP/NPY neurons to ghrelin administration and loss of leptin [25]. In contrast to the situation in BSX^-/-^ knockout animals, AgRP/NPY neuron numbers and expression levels of *Npy* and *Agrp* did not obviously differ between wild-type and *Bsx^Q159K/Q159K^* mice (Suppl. Fig. 3A,B). Furthermore, ghrelin responsiveness, as monitored by *c-Fos* expression, was not affected in BSX^Q159K^-expressing AgRP/NPY neurons (Suppl. Fig. 3A,B). The integrity of the BSX^Q159K^/AgRP/NPY neurons was further tested by generating *Bsx^Q159K/Q159K^/ob/ob* mice. Whereas deletion of *Bsx* in leptin-deficient *ob* mice results in amelioration of hyperphagia and weight gain, as previously reported by us [25], the genetic combination of two *Bsx^Q159K^* alleles with leptin deficiency does not alter the morbid weight gain of *ob/ob* animals (data not shown). In line with this finding, the well-known constitutive activation of AgRP/NPY neurons in the absence of leptin, monitored by *c-Fos* expression, was still present in AgRP/NPY neurons of *Bsx^Q159K/Q159K^/ob/ob* animals (Suppl. Fig. 3C,D) in contrast to what is observed for *Bsx^-/-^/ob/ob* double knockout animals (unpublished observation). These results together strongly suggest that BSX^Q159K^-expressing AgRP/NPY neurons are functionally not impaired. Because *Bsx* expression in the arcuate nucleus is not restricted to AgRP/NPY neurons, we monitored the expression of other hormones in the arcuate nucleus. Unexpectedly, the number of arcuate nucleus *Sst* (somatostatin)-positive neurons was greatly diminished, whereas expression of *Ghrh* (growth hormone-releasing hormone), *Trh* (thyrotropin-releasing hormone) and *Pomc* (proopiomelanocortin) was not affected in the arcuate nucleus of *Bsx^Q159K/Q159K^ mice* (Suppl. Fig. 3A,B and data not shown). These results suggest that the BSX^Q159K^ variant is functional, modulating the outcome of the homeobox code in arcuate nucleus cell-type determination without affecting AgRP/NPY neuron function.

It is important to note here that BSX also affects development of the pineal gland and thereby melatonin levels [38, 39]. However, as C57BL/6J mice are considered “melatonin-deficient” due to a mutation in the gene AANAT (serotonin N-acetyltransferase) [40], we can exclude any melatonin effects to the below described phenotypic differences between wild-type and homozygous BSX^Q159K^ male mice [41].

### Bsx^Q159K/Q159K^ mice show a profound loss of sleep-wake cycles

The compact region of the DMH as well as HDC neurons have both been implicated in the regulation of sleep-wake cycles and arousal [5, 18]. To investigate the consequences of DMHc and HDC neuronal absence in homozygous BSX^Q159K^ animals, we employed EEG/EMG (electroencephalography/electromyography) recording in combination with simultaneous metabolic phenotyping and concurrent infrared (IR) beam-break locomotor activity measurements. Many studies use wheel-running activity to record circadian output. However, it has been shown that wheel-running activity, in contrast to spontaneous locomotor activity (SLA), possesses a motivational component and is strongly reduced in Hdc knockout animals, unlike SLA [42, 43]. Because HDC expressing neurons are absent in Bsx^Q159K/Q159K^ mice SLA measurements were utilized to not distort results.

Although *Bsx^Q159K/Q159K^* female mice have the same anatomical hypothalamic changes as *Bsx^Q159K/Q159K^* male mice, to facilitate comparison with other studies, only male mice were used in all further analyses unless stated otherwise. To be able to record EEG/EMG of unrestrained animals in metabolic cages, we established the NeuroLogger 2A device for C57BL/6J animals, which allows 80 hours of uninterrupted EEG/EMG recording, in combination with the PhenoMaster System, both from TSE Systems/Germany.

Long-term EEG/EMG recordings of homozygous BSX^Q159K^ male mice demonstrated that under 12:12-hour light-dark (LD) conditions, sleep-wake cycles were profoundly lost with an even distribution of wake time, NREM and REM sleep between light and dark period in the absence of any remaining pattern as shown by 60-h hypnograms, whereas overall wake time did not differ between wild-type and *Bsx^Q159K/Q159K^* mice (Fig. 4A and Suppl. Fig. 4A,B). This is important to point out as HDC neurons are absent in homozygous BSX^Q159K^ male mice which have been proposed to govern wakefulness [44]. However our findings are in line with a study that reassessed and questioned the long posited view of a basic wake-promoting role for histaminergic neurons [45].

**Fig. 4:**
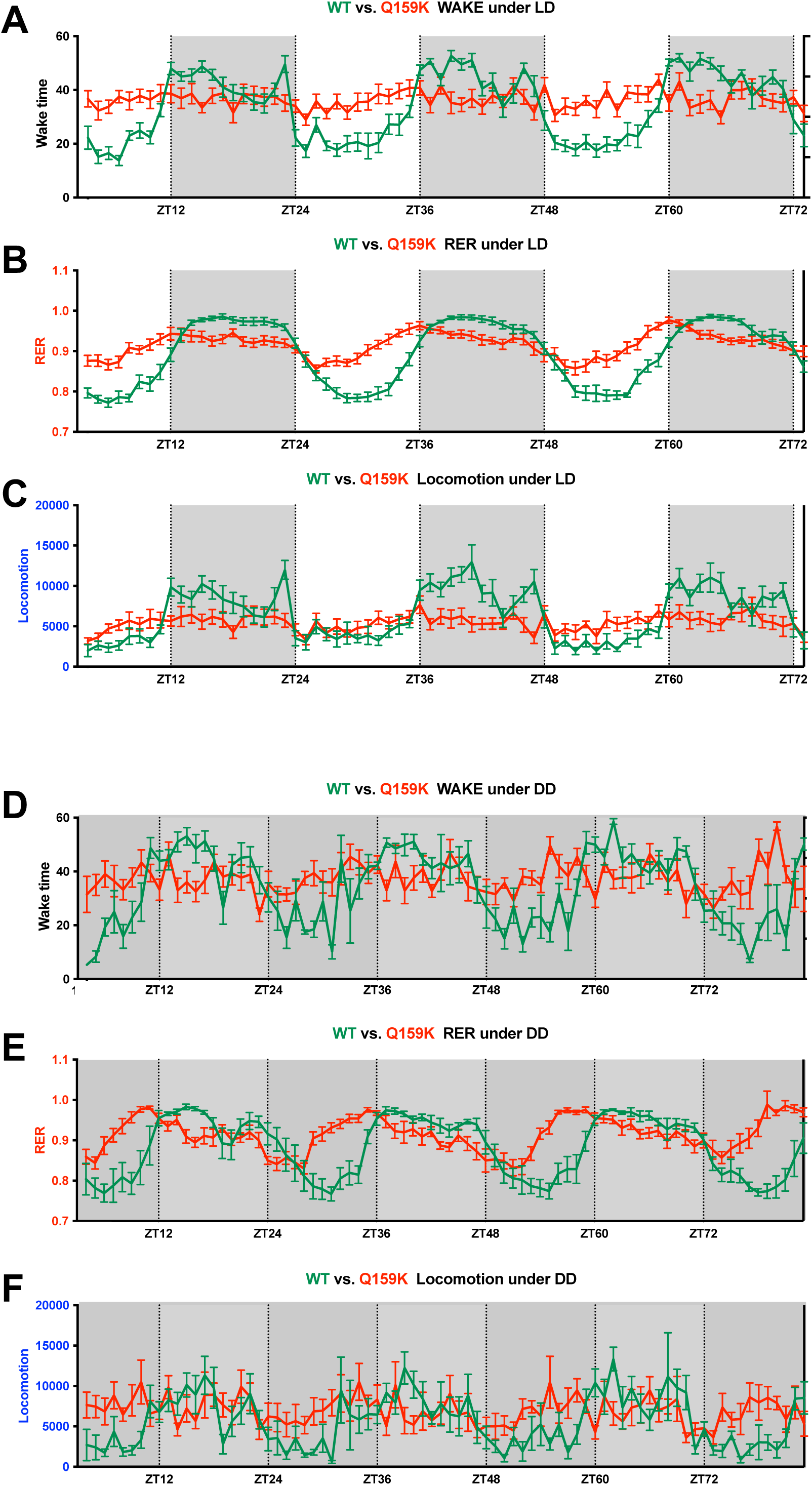
Profound loss of sleep-wake cycles in Bsx^Q159K/Q159K^ male mice. Diagrams are shown as line graphs with mean and error bars (SEM) on the y-axis for wake time (min), respiratory exchange ratio (RER) and locomotion (total beam break counts) respectively and zeitgeber time (ZT) on the x-axis. Grey areas represent time when lights are off. **A)** Wake time of Bsx^WT/WT^ male mice (N=11) cycles as expected according to the circadian 12:12 h light-dark (LD) cycle. In contrast, Bsx^Q159K/Q159K^ male mice (N=15) do not show a difference in wake time between light and dark phase showing more wake time than their Bsx^WT/WT^ littermates during the light phase. **B)** Respiratory exchange ratio (RER) of Bsx^WT/WT^ male mice (N=11) follows the circadian 12:12 h light-dark (LD) cycle with food available ad libitum (higher RER during the active dark phase, indicating greater use of carbohydrates, lower RER during the light phase, reflecting higher fat oxidation) as a result of their circadian feeding behaviour. In contrast, Bsx^Q159K/Q159K^ male mice (N=15) reveal a rhythmic RER pattern that differs substantially from their Bsx^WT/WT^ littermates and does not coincide with the 12:12 dark-light cycle revealing that the daily caloric intake of Bsx^Q159K/Q159K^ male mice has been substantially shifted to the light phase. **C)** Spontaneous locomotor activity of Bsx^WT/WT^ male mice (N=11) follows the circadian 12:12 h light-dark (LD) cycle with elevated locomotor activity as nocturnal animals during the dark phase. In contrast, Bsx^Q159K/Q159K^ male mice (N=15) do not change their spontaneous locomotor activity over a 24 h period. **D)** Wake time of Bsx^WT/WT^ male mice (N=4) cycles in constant darkness (DD). In contrast, Bsx^Q159K/Q159K^ male mice (N=7) do not show a difference in wake time between putative “light” and dark phase being more awake than their Bsx^WT/WT^ littermates during the “light” phase. **E)** Respiratory exchange ratio (RER) of Bsx^WT/WT^ male mice (N=4) displays circadian rhythmicity in DD with food available ad libitum. In contrast, Bsx^Q159K/Q159K^ male mice (N=7) show a robust rhythmic RER pattern that differs substantially and appears to be “phase” shifted under DD relative to their Bsx^WT/WT^ littermates demonstrating a different timely caloric intake between the two genotypes. **F)** Spontaneous locomotor activity of Bsx^WT/WT^ male mice (N=4) follows circadian rhythmicity in DD. In contrast, Bsx^Q159K/Q159K^ male mice (N=7) do not change their spontaneous locomotor activity over a 24 h period under DD setting.

Simultaneous metabolic phenotyping of wild-type and *Bsx^Q159K/Q159K^*mice revealed an attenuated respiratory exchange ratio (RER) rhythm with an ∼24-h pattern reflecting a substantial shift of daily caloric intake from the dark to the light phase of the 12:12-hour LD cycle in *Bsx^Q159K/Q159K^* mice and absence of spontaneous locomotor rhythmicity during 24 h period (Fig. 4B,C). Whereas sleep-wake cycles and locomotor rhythmicity were also profoundly lost in dark-dark (DD) conditions with overall preserved and even distribution of wake time between “light” and dark period in the absence of any remaining pattern as shown by 60-h hypnograms in *Bsx^Q159K/Q159K^* mice, simultaneous metabolic recordings in DD uncovered a rhythmic RER pattern that differs substantially from wild-type mice reflecting an alteration in feeding behaviour (Fig. 4D,E,F and Suppl. Fig. 4C,D). These results show that, in *Bsx^Q159K/Q159K^* male mice, circadian behavioural rhythms are profoundly lost and decoupled from metabolic rhythmicity.

### Bsx^Q159K/Q159K^ mice do not show food-anticipatory activity (FAA)

Whether the DMH is also critical for food-anticipatory activity (FAA) is still debated [10, 11]. To shed light on a possible role of the DMHc in food-anticipatory activity, we entrained wild-type and homozygous BSX^Q159K^ male mice to daytime-restricted feeding with food access for only 4 hours (ZT7–ZT11), during which they had to consume their total daily caloric intake. We only started recording once the animals had adjusted to the new ZT7–ZT11 feeding scheme which was assumed when they had regained their starting body weight. Wild-type animals, as expected, strongly increased their wake time as well as their locomotor activity about 2 hours before food presentation as reported before [46]. In contrast, homozygous BSX^Q159K^ male mice only increased their wake time upon feeding (Fig. 5). These results therefore genetically confirm a critical role of the DMHc as an essential part of the hypothalamic integrator of sleep-wake cycles and food anticipatory activity (FAA), consistent with results obtained by DMH lesion studies [21].

**Fig. 5:**
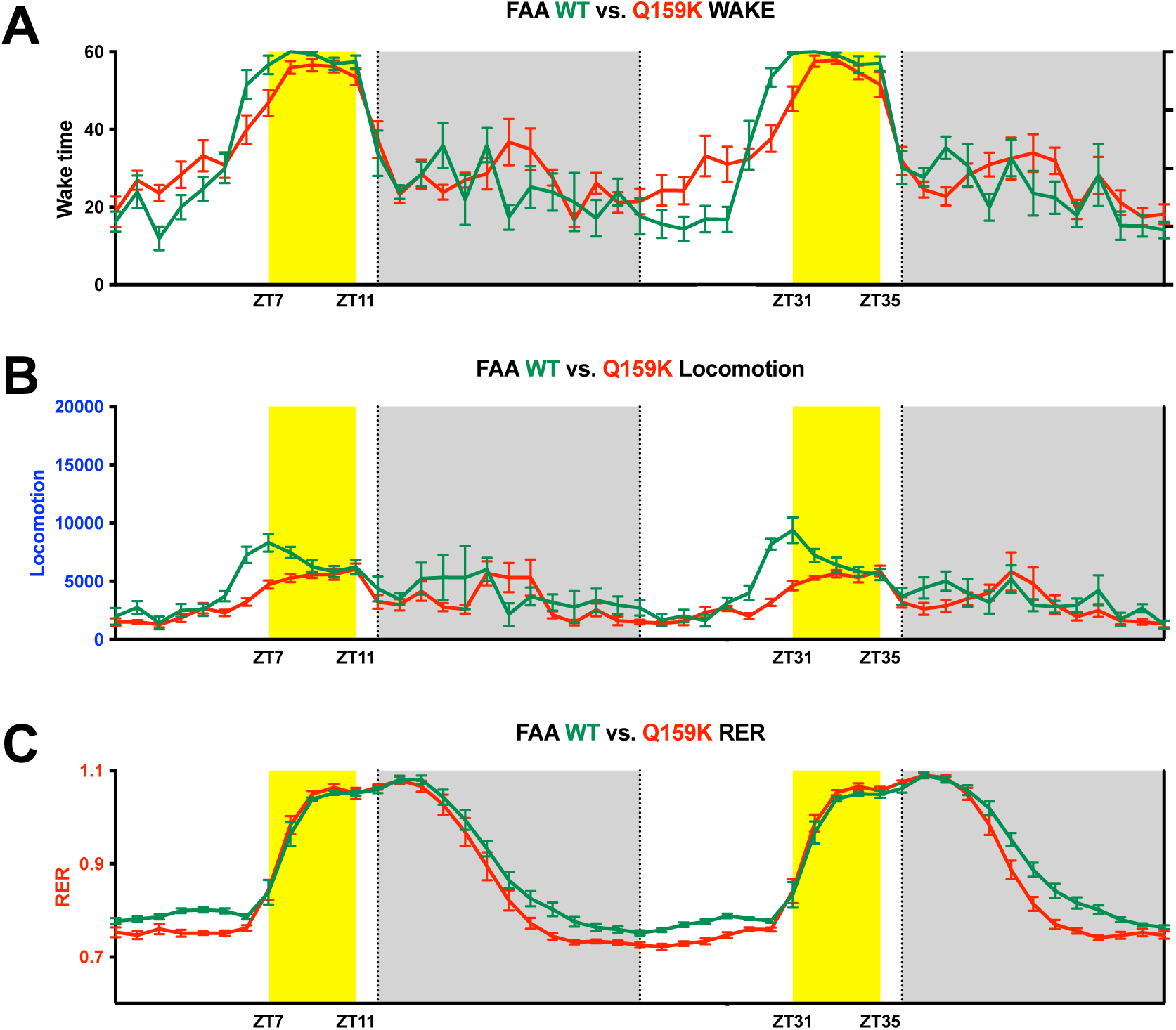
BsxQ159K/Q159K male mice do not show food anticipatory activity (FAA) Diagrams are shown as line graphs with mean and error bars (SEM) on the y-axis for wake time (min), respiratory exchange ratio (RER) and locomotion (total beam break counts) respectively and zeitgeber time (ZT) on the x-axis. Grey areas represent time when lights are off. Yellow areas represent time when food was available. **A)** Bsx^WT/WT^ male mice (N=6) increased their wake time about 2 hours before food became available. In contrast, Bsx^Q159K/Q159K^ male mice (N=8) only increased their wake time upon feeding. **B)** Bsx^WT/WT^ male mice (N=6) concomitantly increased their locomotor activity before food became available. In contrast, Bsx^Q159K/Q159K^ male mice (N=8) barely increased their locomotor activity when food was presented. **C)** Increase in locomotor activity in Bsx^WT/WT^ male mice did not lead to an increase in RER, which was only observed in both genotypes once food was consumed.

### Food-seeking behaviour is attenuated in homozygous BSX^Q159K^ animals

We have previously shown that BSX, in addition to regulating AgRP/NPY expression, is important for locomotory behaviour upon food deprivation [25]. Because, unlike *Bsx^-/-^* knockout animals, AgRP/NPY function is preserved in *Bsx^Q159K/Q159K^* males, we asked whether food-seeking behaviour would also be maintained. Simultaneous EEG/EMG and locomotor recordings during food deprivation (FD) revealed that the locomotor component of the food-seeking response is blunted in *Bsx^Q159K/Q159K^* males. Notably, during the light phase of FD, these mice, like their wild-type littermates, significantly increased wake time, yet their locomotor response remained blunted (Fig. 6A,B). This finding indicates that *Bsx^Q159K/Q159K^*males can detect the metabolic state of deprivation, but do not initiate the expected food-seeking locomotion, consistent with a dissociation between metabolic drive and behavioural output. To determine whether there is a general defect in elevating locomotion, we introduced two standard challenges: novel-cage transfer during the light phase and brief olfactory exposure to food at the end of FD. In both paradigms, *Bsx^Q159K/Q159K^* and wild-type males increased wakefulness and locomotor activity, whereas the locomotor response to fasting itself remained absent in *Bsx^Q159K/Q159K^* male mice during the time between these challenges (Suppl. Fig. 5A,B). Together, these data indicate a context-dependent deficit in food-seeking type locomotion rather than a global reduction in the capacity to increase locomotor activity.

**Fig. 6:**
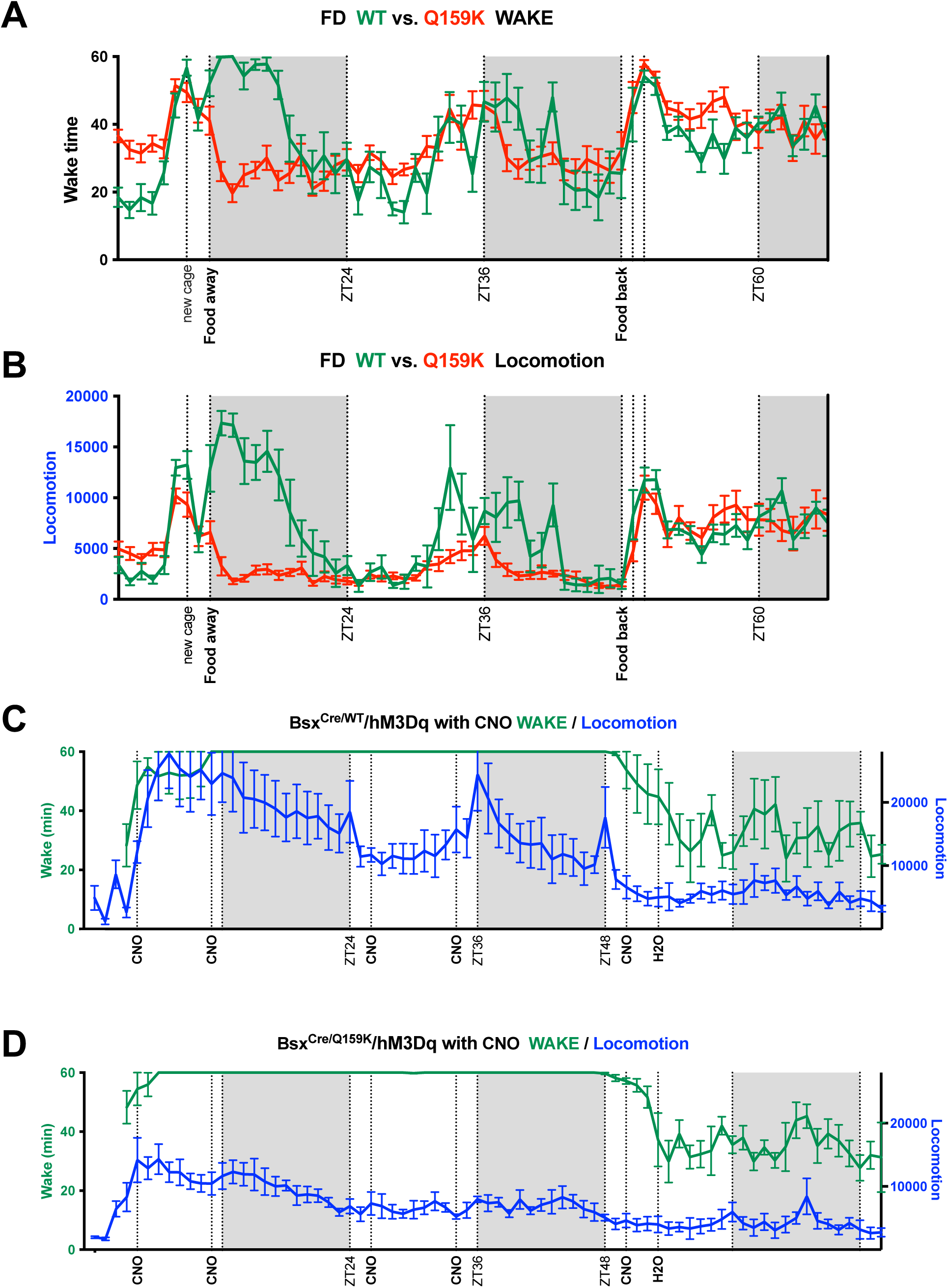
BSX network activity is required and sufficient for food-seeking behaviour. Diagrams are shown as line graphs with mean and error bars (SEM) on the y-axis for wake time (min) and locomotion (total beam break counts) respectively and zeitgeber time (ZT) on the x-axis. Grey areas represent time when lights are off. **A)** Acute prolonged food deprivation (FD) leads to increase of wake time during the light phase of FD in Bsx^WT/WT^ male mice (N=8) and Bsx^Q159K/Q159K^ male mice (N=12) demonstrating that both genotypes are able to sense the metabolic state of food deprivation. **B)** Acute prolonged food deprivation leads to strong increase of locomotor activity during the light phase in Bsx^WT/WT^ male mice (N=8) but not in Bsx^Q159K/Q159K^ male mice (N=12) demonstrating decoupling of metabolic state sensing and behavioural execution in Bsx^Q159K/Q159K^ male mice. **C)** Application of CNO (0.5 mg/ml) through the drinking water to male mice that express hM3Dq in neurons of BSX^WT^ lineage (N=4) maintain >36 h of wakefulness with concomitant elevation of locomotor activity throughout the whole CNO application period. **D)** Application of CNO (0.5 mg/ml) through the drinking water to male mice that express hM3Dq in neurons of BSX^Q159K^ lineage (n=5) induce >36 h of wakefulness without elevated locomotor activity during the light phase despite prolonged wakefulness over the CNO application period.

These results show that HDC neurons are not required for basal wakefulness and that certain forms of locomotor activity do not depend on their presence. To assess whether HDC neurons contribute to prolonged wakefulness, we took advantage of the following observation: during c-Fos whole brain mapping after BSX network activation by a single CNO injection, *Bsx^Cre/WT^/Rosa-hM3Dq* mice displayed a marked increase in exploratory locomotion and EEG/EMG-defined prolonged wakefulness for ∼3 h, accompanied by increased water and food intake (data not shown). Prolonged wakefulness elicited by BSX-network activation was accompanied by activation of paraventricular nucleus (PVN) corticotropin-releasing hormone (CRH) neurons and lateral hypothalamic (LH) Orexin/Hypocretin neurons as demonstrated by strong *c-Fos* expression in both neuronal populations which do not express *Bsx* themselves (Suppl. Fig. S6) [47–50]. This prompted a self-administration paradigm, in which CNO was added to the drinking water, to create a self-reinforcing cycle of CNO application. Upon application of CNO through the drinking water, *Bsx^Cre/WT^/Rosa-hM3Dq* mice maintained >36 h of wakefulness and elevated locomotor activity, which ceased when CNO was withdrawn (Fig. 6C,D and Suppl. Fig. 5C,D). These results demonstrate that the BSX-network is sufficient to elicit prolonged food-seeking behaviour. In contrast, *Bsx^Cre/Q159K^/Rosa-hM3Dq* mice also maintained >36 h of wakefulness with activation of CRH and orexin neurons upon CNO application, but their locomotor activity was barely elevated when compared with *Bsx^Cre/WT^/Rosa-hM3Dq* mice (Fig.6C,D). Thus, HDC neurons and DMHc are not required to maintain prolonged wakefulness at least when elicited through BSX-network activation, whereas HDC neurons, alone or together with the DMHc, appear important for amplifying locomotor activity/food-seeking distance during negative energy balance [51]. We note that locomotor activity is an unreliable proxy for wakefulness in this context and caution is warranted when inferring the wake state from locomotor readouts alone.

### Aged homozygous BSX^Q159K^ mice maintain a lean body composition

The ability of an organism to adjust to circadian changes influences ageing: circadian misalignment between internal clocks and daily cycles can impair tissue homeostasis, sleep regulation, and behaviour, whereas high-amplitude circadian rhythms have been associated with longer lifespan in rodents [52, 53]. Mouse lines deficient in clock genes, including *Clock^−/−^* and *Bmal1^−/−^*, develop metabolic disease and show shortened lifespan, whereas *Cry1/Cry2* double-knockout mice are leaner than their wild-type littermates [54–57]. Because Bsx^Q159K/Q159K^ animals show profound loss of circadian behavioural rhythms (sleep-wake cycle, locomotor activity, and feeding behaviour) under light-dark conditions, we examined their lifespan.

We aged C57BL/6J wild-type and *Bsx^Q159K/Q159K^* male littermates derived from *Bsx^WT/Q159K^* intercrosses to determine survival under standard husbandry (12:12h light-dark; ad libitum food and water). Both *Bsx^Q159K/Q159K^* males and their wild-type siblings lived up to 1,000 days, which is within the expected maximal lifespan reported for C57BL/6J mice under comparable conditions [58], with no difference in median or maximum lifespan between genotypes observed (Fig. 7C). In contrast, body composition diverged with age: *Bsx^Q159K/Q159K^*males maintained stable body weight from 3 months onward, whereas wild-type littermates showed the expected age-dependent increase in fat mass. At 80-weeks of age, *Bsx^Q159K/Q159K^* males had ∼20% body fat compared with ∼40% in wild-type littermates (Fig. 7A,B and Suppl. Fig. 7). Although neither genotype exhibited severe health deficits at 110-weeks of age, coat greying was observed only in wild-type animals at this time (Fig. 7D). Taken together, these data indicate that loss of circadian behavioural rhythms in *BSX^Q159K/Q159K^* males does not shorten lifespan under food ad libitum, standard laboratory conditions and is even associated with a leaner body composition. Within the limits of the endpoints measured (body composition, survival, and general well-being), *Bsx^Q159K/Q159K^* males display healthspan and lifespan comparable to wild-type littermates.

**Fig. 7:**
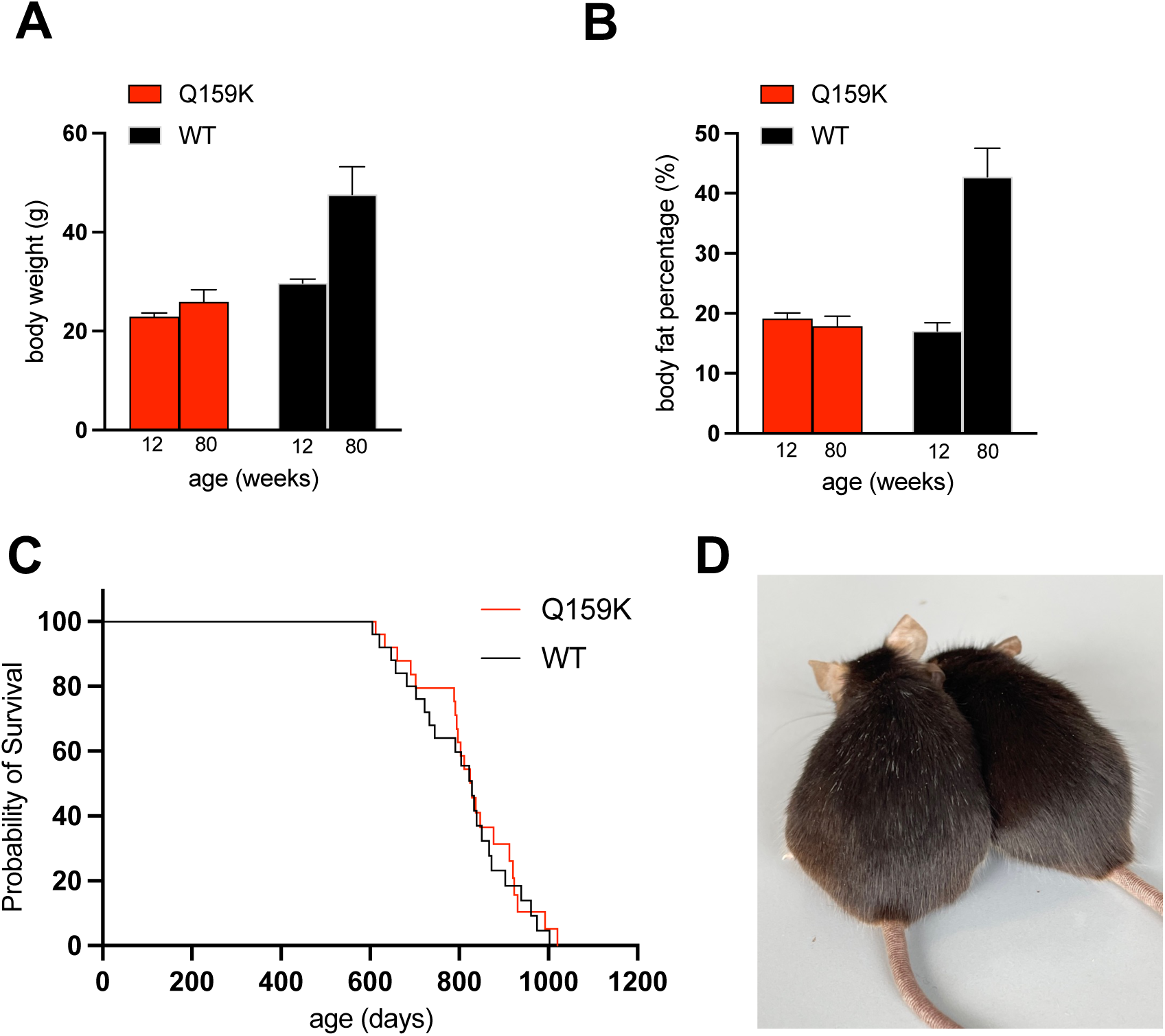
Aged Bsx^Q159K/Q159K^ mice maintain a lean body composition. **A)** Body weight of Bsx^WT/WT^ male mice at 12 week (N=15) and 80 weeks (N=14) and Bsx^Q159K/Q159K^ male mice at 12 week (N=14) and 80 weeks (N=13) showing that only Bsx^WT/WT^ but not Bsx^Q159K/Q159K^ mice substantially gain weight while ageing. Error bars indicate mean and SD. **B)** NMR measurements of body fat percentage of Bsx^WT/WT^ and Bsx^Q159K/Q159K^ male mice reveals that while body fat percentage nearly doubles in Bsx^WT/WT^ mice it remains unchanged in Bsx^Q159K/Q159K^ mice during ageing. Columns in A) and B) represent the same animals, respectively. Error bars indicate mean and SD. See also Suppl. Fig.7. **C)** Kaplan-Meier survival curves for Bsx^WT/WT^ (N=23) and Bsx^Q159K/Q159K^ (N=22) male mice are not significant different between the two genotypes (Unpaired t test, two-tailed P-value: ns) **D)** Representative picture of Bsx^WT/WT^ and Bsx^Q159K/Q159K^ littermate male mice at 110 weeks of age reveals only hair greying in wild-type animal.

## Discussion

Classical work on light-entrained (SCN-driven) circadian behavioural rhythms, through SCN lesions or disruption of core clock genes, has yielded a detailed molecular framework linking circadian timing to metabolism and lifespan [1, 2, 6, 59]. Parallel work on the organisation of food-entrainable rhythms, and the question whether a single food-entrainable oscillator (FEO) exists outside the SCN or instead emerges from a distributed network, still remains open [10, 11] . The dorsomedial hypothalamus (DMH) has been implicated in integrating temporal and metabolic cues, thereby shaping circadian rhythms, but its precise role is still evolving [8, 12, 59–61].

Our study provides genetic evidence that the compact region of the dorsomedial hypothalamus (DMHc) and the histaminergic system of the tuberomammillary nucleus (TMN) are required for normal circadian behavioural rhythms. In *Bsx^Q159K/Q159^*^K^ mice, DMHc is anatomically and molecularly absent, as demonstrated by the loss of the characteristic “butterfly-shaped” gene expression pattern of DMHc markers including *Ppp1r17*, *Grp* and *Cck*. In addition, TMN histaminergic neurons are not present, as demonstrated by the absence of their key transcriptional signature genes: *Hdc*, *Prph*, *Ox2r*, and *Wif1* [35] which has also been observed in zebrafish *Bsx* mutants [62].

Consistent with these anatomical changes, we observe key behavioural changes in *BSX^Q159K/Q159K^* mice. First, *Bsx^Q159K/Q159K^* mice have a profound loss of circadian behavioural rhythms, including sleep-wake cycle and the day-night organisation of locomotor activity and feeding, under both LD and DD. Second, *Bsx^Q159K/Q159K^* mice lack food-anticipatory activity (FAA) during daytime-restricted feeding, yet still consume their full daily intake within the 4-h window, demonstrating a dissociation between anticipatory arousal and caloric consumtion. These two key behavioural changes, based on a single amino acid substitution, provide strong genetic support for lesion studies that reached the same conclusion that the DMH is crucial for sleep-wake cycles and FAA [21, 63]. Although our study shows a critical role for the DMHc in FAA, we acknowledge that our data do not prove that the DMHc contains the FEO. Third, metabolic rhythmicity persists in *Bsx^Q159K/Q159K^* mice, as the respiratory exchange ratio (RER) retains a detectable ∼24 h rhythm, which is more pronounced under DD and attenuated under LD.

We note that these changes occur in the presence of a most likely normally cycling SCN under LD, based on the fact that (1) BSX is not expressed in cells in or around the SCN (2) ISH gene expression snapshots of the SCN at ZT7 in *Bsx^WT/WT^* mice and *Bsx^Q159K/Q159K^* mice do not show any difference between genotypes and reveal optically dense expression of *Per2* [64, 65] (Suppl. Fig. 8), which shows that gene expression of the SCN in both genotypes corresponds to the expected ZT7 gene expression pattern, despite overall wake time of *Bsx^Q159K/Q159K^* mice during the light phase is significantly higher than that of their wild-type littermates (Suppl. Fig. 4A).

The metabolic data further provide insight how behaviour and metabolism decouple. In *Bsx^Q159K/Q159K^* mice, an RER rhythm persists despite loss of behavioural cycling, indicating that metabolic oscillators can operate when behavioural timing is disturbed.

In *Bsx^Q159K/Q159K^* mice, AgRP/NPY neuron numbers and neuropeptide expression are comparable to wild-type littermates, with ghrelin-evoked *c-Fos* expression still being intact. Furthermore, leptin deficiency in *Bsx^Q159K/Q159K^* mice still produces morbid weight gain in contrast when BSX is absent. Taken together, these findings indicate that the hunger-driving AgRP/NPY neurons and the satiety brake neurons in the PVN remain operational, whereas DMHc neurons, which regulate sleep-wake cycles and thus act as a temporal timer, are missing, which may contribute to the observed change in feeding behaviour across the 24 h cycle in *Bsx^Q159K/Q159K^* mice. How much of this RER rhythm in the absence of the DMHc in Bsx^Q159K/Q159K^ mice is driven by AgRP/NPY neurons, which have been implicated in the circadian control of daily feeding rhythms or the liver clock that has been shown to be able to cycle independently of SCN input, or both together through their interconnectedness, remains to be seen [22, 60, 66–68].

In *Bsx^Q159K/Q159K^* mice that lack TMN histaminergic neurons, the reduced RER amplitude in LD compared with DD is consistent with weaker coupling of photic input to arousal and feeding. By contrast, in DD, a robust rhythmic RER pattern is evident when neither light nor histaminergic drive is present [69–71]. We acknowledge that this interpretation aligns with the observed pattern rather than providing a direct demonstration.

Our genetic manipulations also clarify how these co-affected nodes influence arousal and locomotion. With chemogenetic activation of the BSX network, both genotypes can sustain >36 h of wakefulness, indicating that TMN histaminergic neurons are not required to maintain wakefulness per se. However, locomotor activity differs: wild-type mice show marked increases in locomotory movement, whereas *BSX^Q159K/Q159K^* mice display prolonged wakefulness with little or no locomotor activity increase [48, 72, 73]. Moreover, food-anticipatory activity locomotion that precedes restricted feeding and elevated locomotion that accompanies food deprivation is blunted in *Bsx^Q159K/Q159K^* mice, particularly during the light phase, even though these same mice can increase locomotion when placed in a new cage or exposed to food odour. These behavioural patterns are consistent with a role for the DMHc in scheduling/anticipation and for the histaminergic system in context-dependent locomotor activity modulation, while acknowledging that the present data cannot asign effects definitively between the two.

We have shown that loss of *Bsx* in leptin-deficient *ob/ob* mice ameliorates their morbid obese phenotype due to AgRP/NPY neuron impairment. In contrast, loss of DMHc neurons in *Bsx^Q159K/Q159K^/ob/ob* mice does not prevent obesity development, which is in line with prior findings that reported that chemogenetic activation of DMH PPP1R17 neurons in *ob/ob* mice decreased food intake and body weight, whereas inhibition increased both [16]. Independently, another study similar demonstrated that chemogenetic activation of a refeeding-activated DMHc neuronal population that expresses *Cck* and *Grp* also leads to decrease in food intake [15]. If the two studies describe the same or independent DMHc populations remains to be seen. Regardless, as *Ppp1r17*, *Cck* and *Grp* markers are absent in our model from DMHc in *Bsx^Q159K/Q159K^* mice, consistent with *in situ* data, the DMHc “brake” on food intake is most likely diminished. In contrast to *Bsx^Q159K/Q159K^* mice, which remain lean at advanced age, it has been shown that DMH-specific *Ppp1r17* or *Prdm13* knockdown increased fat mass during ageing in mice [17, 74]. Whether, and how, missing DMHc CCK- or GRP-positive but PPP1R17-negative neurons in *Bsx^Q159K/Q159K^* mice contribute to this different outcome remains to be determined. Regardless, our results further strengthen a role for the DMHc as part of a hypothalamic-adipose communication axis [17]. We note that TRH neuron number and distribution in the DMH, which have been shown to relay information from the SCN to AgRP/NPY neurons, are not affected in *Bsx^Q159K/Q159K^*mice [66]. Thus, our findings contribute to and further strengthen a role for DMHc cell types in regulating feeding behaviour and metabolism.

At the level of arcuate nucleus cell types, BSX^Q159K^ notably remodels the arcuate nucleus while preserving AgRP/NPY neurons, in contrast to when BSX is deleted. Arcuate SST-positive neurons are, however, reduced whereas GHRH neuron numbers are not affected. In global somatostatin knockout mice, circulating growth hormone (GH) is increased, yet IGF-1 and body size do not rise [75]. Why *Bsx^Q159K/Q159K^* mice have smaller body size despite fewer SST-positive neurons remains unclear; one speculative possibility is that loss of sleep-wake cycling disrupts GH rhythmicity, given the well-established coupling between sleep and GH release; however, direct endocrine measurements in our model have been inconclusive. By contrast, *Pomc* and *Trh* expression in the arcuate nucleus are not altered. This selective hormonal cell-type pattern in the arcuate nucleus, together with the absence of the DMHc and TMN histaminergic neurons, suggests that the Q159K missense variant perturbs BSX-dependent developmental specification programmes. While direct BSX targets remain to be identified, the data support the view that arcuate, DMHc, and TMN histaminergic lineages share BSX-dependent developmental requirements that are altered by the BSX^Q159K^ AA substitution.

Finally, the long-term phenotypes place these rhythm changes in physiological context. *Bsx^Q159K/Q159K^* males remain lean with stable body weight during ageing, and lifespan is not different from wild type littermates under standard husbandry (12:12 LD; ad libitum food). Within the limits of the endpoints measured (body composition, survival, and general well-being), these data indicate that overall metabolic balance is preserved despite loss of circadian behavioural rhythms, while the temporal structure of metabolism (e.g., RER rhythm under LD) is altered. Further metabolic challenge studies in *Bsx^Q159K/Q159K^* mice are warranted to determine how metabolism that is not constrained by behavioural rhythms, yet retains metabolic oscillations, responds to temporal (e.g., restricted feeding, altered photoperiod) and energetic (e.g., caloric restriction, high-fat diet) alterations.

In summary, the *Bsx^Q159K/Q159K^* model demonstrates that removing the DMHc together with the TMN histaminergic system abolishes sleep-wake cycles and FAA and disrupts broader circadian behavioural rhythms, while sparing a residual metabolic rhythm and the core AgRP/NPY neuron-to-PVN axis that drives feeding. These findings support the view that circadian behavioural rhythmicity is a context-dependent adaptation whose principal advantages emerge under diurnal conditions. Therefore, under continuous food availability, metabolic homeostasis and body composition may be maintained, or even improved, when circadian behaviour and metabolism are not tightly coupled by a central hypothalamic circadian integrator [12].

## Methods

### Animals

All animal procedures including generations of transgenic mice were approved by competent authority, the Berlin State Office for Health and Social Affairs (LAGeSo), Germany. All animals were group-housed under a controlled 12:12h Light : Dark (LD) cycle, room temperature 20-25 °C, humidity level of 45% to 65% and fed 2019S Teklad Global 19% Protein Extruded Rodent Diet with chow and water available *ad libitum*.

The *Bsx^Q159K^* allele was generated by CRISPR/Cas9 genome editing in C57BL/6J (JAX® mice) from Charles River/Germany, using a targeting guide RNA (gRNA: GCT TTT TAT GCT TCA TCC GC) and a 120bp oligo (CCG CTT TCG GTT CGT CTT GGC TTT TCC TCA GCT GCT TTT TAT GCT TCA TCC GCC GAT TCT TGA ACC ACG TTT TCA CCT GAT GTC GGG GAG ACA AAC AAG ATC ATA TCA TTC CTC ACA CAA). Guide RNAs and Oligos for all other reported lines carrying BSX amino acid missense variants can be obtained upon request. In all analysis/experiments wild-type and *Bsx^Q159K/Q159K^* male littermates derived from *Bsx^WT/Q159K^*intercrosses were used if not stated otherwise. The *Bsx^CRE^* knock-in allele was generated by homologous recombination in R1 (129S1 x 129X1) ES cells by placing Cre recombinase at the ATG start site of the *Bsx* gene which at the same time resulted in a BSX loss-of-function allele (exact sequence of cloning vector available upon request). After injection of ES cells into C57BL/6J blastocysts, positive animals were backcrossed at least eight generations to C57BL/6J mice. The Jackson Laboratory strains: B6 *ob* (#000632), Rosa-Ai14-tdTomato (#007914) and R26-hM3Dq/mCitrine (#026220) were backcrossed on C57BL/6J if required.

### Histology

*Vibratome sectioning on Leica VT1200S*: 2% PFA fixed brains were embedded in 2% agarose in PBS. When the agarose solidified the brains were trimmed and placed on the vibratome holder using cyanoacrylate glue. Then, the vibratome chamber was filled with cold PBS. 50 μm sections were cut at a blade frequency of 9 Hz and speed of 0,5 mm/s. The remaining agarose attached to the sections was carefully removed using fine brush and forceps. The collected sections were either processed immediately or stored in PBS at 4 °C. *Cryotome sectioning on Leica CM3050S*: 10% Formalin fixed brains were embedded in Leica Cryo-Gel medium and frozen in a dry ice/ethanol bath. 16 μm sections were cut and collected on Superfrost Plus Gold slides. The collected sections were either processed immediately or stored at -80 °C.

*Immunohistochemistry:* 50 μm free-floating brain sections were used for immunofluorescence staining. First, the sections were incubated in blocking buffer (3 % normal donkey serum or 3 % normal goat serum, 3 % BSA, 0.3 % Triton X-100 in PBS) for 1 hour at RT. Then, the sections were incubated with primary antibody diluted in 1 % BSA, 0.3 % Triton X- 100 in PBS at 4 °C overnight. The following day, the sections were washed three times with the 0.3 % Triton X-100 in PBS for 20 min at RT. Next, the sections were incubated for 2 hours with a 1:700 dilution of the secondary antibody diluted in 1 % BSA, 0.3 % Triton X-100 in PBS at RT. Subsequently, the sections were washed three times for 20 min with the washing buffer (0.3 % Triton X-100 in PBS) at RT. Finally, the brain sections were mounted on slides, embedded in ProLong Diamond, cover-slipped, and analyzed using a Leica TCS SP5 confocal microscope. The primary and secondary antibodies were used at the following concentrations: rabbit anti-BSX (1:200). Secondary antibodies: donkey anti-rabbit IgG Alexa488 (1:700) (Jackson ImmunoResearch) and goat anti-rabbit IgG Alexa594 (1:700) (Life Technologies). All histological staining pictures show representative sections.

*In situ hybridization:* 16 μm cryosections were used for in situ hybridization with RNAscope following the manufacturer’s instructions and protocol. The following RNAscope probes were used: RNAscope Probe-Mm-Agrp #400711, RNAscope Probe-Mm-Bsx #538371, RNAscope Probe-Mm-Cck#402271, RNAscope Probe-Mm-Fos#316921, RNAscope Probe-Mm-Ghrh#470991, RNAscope Probe-Mm-Grp#317861, RNAscope Probe-Mm-Hcrt #490461, RNAscope Probe-MmHcrtr2#460881, RNAscope Probe-Mm-Hdc#490471, RNAscope Probe-Mm-Lepr#402731, RNAscope Probe-Mm-Nms#472331, RNAscope Probe-Mm-Npy#313321, RNAscope Probe-Mm-Per1#438751, RNAscope Probe-Mm-Per2#454521, RNAscope Probe-Mm-Pomc-O1#519151, RNAscope Probe-Mm-Ppp1r17#497351, RNAscope Probe-Mm-Prph#400361, RNAscope Probe-Mm-Sst-01#482691, RNAscope Probe-Mm-Trh#436811, RNAscope Probe-td Tomato#317041, RNAscope Probe-Mm-Wif1#412361. All *in situ* hybridization pictures show representative sections.

*Light-sheet imaging:* Light-sheet imaging and c-Fos mapping was done together with LifeCanvas Technologies U.S.A.: Briefly, CNO activated and paraformaldehyde-fixed brains from BSX^Cre/WT^/Ai14-tdTomato/R26-hM3Dq or BSX^Cre/Q159K^/Ai14-tdTomato/R26-hM3Dq genotypes were immunostained with c-Fos/NeuN antibodies, cleared and subsequently imaged at LifeCanvas Technologies.

### Image Analysis and Visualization Workflow

The best two representative whole-brain datasets with three fluorescence channels (488 nm NeuN, 647 nm cFOS, 561 nm tdTomato) were processed. First, raw TIFF stacks are converted to OME-Zarr with defined voxel size and chunking to enable efficient, multi-resolution access. The reference atlas is also upsampled and written to OME-Zarr. Each dataset is then nonlinearly warped into atlas space using precomputed registration parameters provided by the experts at LifeCanvas who acquired the image data. From the atlas-space data, we render slice series along the coronal, sagittal, and transverse planes within a predefined crop region of the Hypothalamus and render atlas label borders for the same planes. These are encoded into per-dataset/per-channel videos, blended in pairs and rendered side-by-side for comparative video exploration. To quantify regional signals, we create atlas-masked maximum-intensity projections for selected target regions (e.g., DMH, PH, TU, TM), computing per-dataset/channel metrics that are aggregated and plotted. All scripts are provided through a reproducible NextFlow – workflow which can be obtained upon request.

### Electroencephalography (EEG)

*EEG surgical procedure and analysis*: Surgery for sleep-wake cycle measurements was performed on mice older than 12 weeks. Before surgery mice were injected subcutaneously with Carprofen as pain treatment (5 mg/kg bodyweight). Male mice were anesthetized using isoflurane (induction, 5 %; maintenance, 2%, flow rate 200 ml/min) in oxygen-enriched air and head fixed on a stereotaxic frame (Model 940, David Kopf Instruments). Body temperature was maintained at 37 °C using a heating pad. Eyes were protected with Bepanthen ointment and scalp was shaved and cleaned with ethanol before making a longitudinal incision along the midline to reveal skull and neck muscle. Skull periosteum was removed and thoroughly cleaned with ethanol. A carbide cutter was used to drill 5 holes into the skull with the coordinates: frontal bone (-1 mm to Bregma, ± 1.5 mm from midline), parietal bone (-1 mm to Lambda, ± 1.5 mm from midline) and interparietal bone (+2 mm from Lambda, 0.0 mm from midline). Gold-plated EEG electrodes each with a connector cable and a metal pin were implanted through the skull onto the dura mater using a screw driver: 2 EEG recording electrodes at frontal bone, 2 EEG recording electrodes at parietal bone and 1 reference electrode at interparietal bone. Teflon-insulated stainless-steel wire was inserted in the neck muscle to record the EMG signal and the skin near the neck was sewed by surgical suture. The skull was covered with a thin layer of dental self-adhesive and dual-curing high-performance composite cement (Rely Unicem 2 Automix, 3M) and light cured using a LED Curing Light (Elipar DeepCure-S, 3M). The EEG electrodes and the EMG wire were connected into an adaptor. Finally, dental cement was applied to cover all the electrodes and wires completely. After recovery in a quiet and warm area, mice were housed singly in their home cage and Carprofen was administered daily for 3 days. After 2 weeks of recovery the mice were acclimatized to a mock logger (similar size and weight to the NeuroLogger 2A from TSE Systems) for 2 days before the experiment. For natural sleep recordings animals were connected to the NeuroLogger 2A and mice were left undisturbed for at least 80 hours to enable three full sleep-wake cycles (12:12h light-dark) (LD) or constant darkness (DD) to be recorded.

*EEG data acquisition, processing and the sleep-wake state classification:* The EEG and EMG signals were digitalized by the NeuroLogger 2A with the sampling rate of 500 Hz. Original data acquired in hexadecimal format (.hex) were converted into European data format (.edf). Sleep states were scored with the sleep analysis software (SleepSign, Kissei Comtec)[76]. EEG and EMG signals were filtered by band-pass between 0.20-100 Hz and 30-300 Hz, respectively. The SleepSign software calculated the power spectrum of EEG waveform of every 5-s epoch by Fast Fourier transform (FFT) algorithms (FFT points: 1024, the average of 5 spectra per epoch). All scoring was screened automatically based on of the signature of EEG and EMG waveforms in 5-s epochs. The general criteria for classifications of the different states were: Wake = with locomotor activity, high EMG, and intermediate theta/delta wave ratio; NREM sleep = no locomotor activity, low EMG, and predominated with delta wave; REM sleep = no locomotor activity, low EMG, and predominated with theta wave. All classifications of states automatically assigned by SleepSign were examined visually and corrected manually. The total durations of wake, NREM and REM at every hour and every 12 hours were exported as a text file.

### Metabolic Phenotyping

Metabolic phenotyping was performed using the PhenoMaster system from TSE Systems. This equipment allows the simultaneous measurement of indirect calorimetry, locomotion, temperature and food intake of singly housed mice. The analysis cages are located in climate chambers with controlled temperature, light and humidity. Before the experiment the mice were trained for 5 days, this training period allowed the mice to learn how to eat and drink from the special feeders. Then, the animals were acclimatized to the metabolic cages with single housing for 4 days. After the training and acclimatization phase, the mice were measured during five consecutive days in the metabolic cages at constant 22 °C, 55% humidity and a 12:12-h light-dark cycle LD or in 24-h DD. During this time, data for energy expenditure, RER, spontaneous locomotor activity, food and water intake and EEG/EMG was collected if mice carried a NeuroLogger 2A.

*Indirect calorimetry:* Energy expenditure (EE) and Respiratory Exchange Ratio (RER) was determined using indirect calorimetry. The PhenoMaster system sampled for 120 seconds every 10-15 minutes the levels of oxygen (O2) and carbon dioxide (CO2) of each metabolic cage. VO2 and VCO2 were calculated as the difference between the reference cage and the sample cage. The resulting energy expenditure was then normalized to the lean mass of each animal. RER was calculated as the ratio of VCO2 and VO2. For calculation of energy expenditure and RER the acquired values for each hour were averaged.

*Locomotor activity measurements:* Spontaneous cage activity was recorded using an infrared frame positioned outside of the metabolic cage in the x- and y-axis. The scan rate of the infrared beans was 100 Hz and the recording interval was 1 minute. Spontaneous ambulatory activity was displayed as the summation of all beam breaks during one hour

*Food and water intake measurements:* Food and water intake was continuously measured during the experiments via weight sensors positioned above the feeders. The cumulative food or water intake for each hour was calculated.

*Food deprivation experiment:* Food-seeking activity and rebound feeding were analyzed using the PhenoMaster system in combination with NeuroLogger recordings. For this purpose, mice were acclimatized to the metabolic cages for 5 days. Next, the basal metabolic parameters (energy expenditure, RER, locomotor activity, food and water intake) and EEG/EMG were recorded at 22 °C, 55 % humidity and a normal 12:12-hour light- dark cycle for 3 days. At the beginning of the dark phase in the third day food was taken away. The food deprivation lasted maximum 36 hours, during this time all previously mentioned metabolic parameters were recorded. Latest after 36 hours food was given back and metabolic parameters of the animals were recorded for at least another 24 hours. For the food smell experiment, food/chow was placed on top of the air intake slot of the metabolic cages unreachable for the mice. In all food deprivation experiments it was secured that no food/chow remained in the experimentation room during food deprivation.

*Food anticipatory activity (FAA):* Food anticipatory activity was analyzed using the PhenoMaster system in combination with NeuroLogger recordings. For this purpose, mice were acclimatized to the metabolic cages for 5 days. Next, the basal metabolic parameters (energy expenditure, RER, locomotor activity, food and water intake) and EEG/EMG were recorded at 22 °C, 55 % humidity and a normal 12:12-hour light- dark cycle for 3 days. Food was then gradually restricted so that finally food was available only during ZT7-ZT11 within a normal 12:12-hour light-dark cycle. Recordings were not started before animals had adjusted to the restricted feeding scheme which was assumed when animals had regained their starting body weight. In all food anticipatory activity experiments it was secured that no food/chow remained in the experimentation room when food was not available.

### Designer Receptors Exclusively Activated by Designer Drugs (DREADDs)

The clozapine-N-oxide (CNO) was sourced from Tocris Biosience (#4936). A working solution containing 0.5 mg/ml in 0.9 % saline was prepared. Either mice were i.p. injected with 5 mg/kg CNO or received CNO (0.5 mg/ml) through the drinking water with 5mM saccharin. DREADD experiments were performed in the metabolic phenotyping system (PhenoMaster) from TSE Systems either with or without recording of the EEG/EMG with NeuroLogger 2A from TSE Systems.

#### Ghrelin response test

Ghrelin induced c-Fos expression in AgRP/NPY was tested in 10-12 weeks old BSX^Q159K/Q159K^ male mice. On treatment day, mice were intraperitoneally injected with 500µl of ghrelin (0.8g/ml dissolved in saline) in the early light phase. Ghrelin was sourced from Biotechne #1459. Mice were sacrificed 90 minutes after injection and perfused transcardially with 10% formaldehyde; brains were removed and post-fixed over night at 4°C as described previously for *in situ* hybridisation.

### Body weight and composition measurements

Mice were weighed and analyzed between 9 and 11 a.m. All animals were ad libitum fed before measurements. Body weight was measured using a standard laboratory scale. Body composition was determined using Nuclear Magnetic Resonance (NMR) equipment from EchoMRI, Houston, USA. For this purpose, non-anesthetized mice were introduced into mouse holder tubes. The tubes were inserted in the MRI equipment and the body composition analysis was performed. The calculated lean mass and fat mass was stored.

### Statistical analysis

Values were expressed as mean ± SEM or ±SD. Data were analyzed using GraphPad Prism 9.0. Data were analyzed using unpaired t test. In all statistical test, data were considered to be significant if p < 0.05.

## Author contributions

Conceptualization: AC.T., C.K., IS.F.; M.T.; Methodology: F.B., J.G., C.K., M.K., FR.R., D.S.,

AC.T. Investigation: F.B., J.G., C.K., M.K., FR.R., D.S., AC.T., C.K., and M.T. Supervision: AC.T., C.K., D.S. and M.T. Writing: M.T.

## Acknowledgments

We are grateful to the staff of the animal facility of the Max-Delbrück Center for animal husbandry, in particular Sina Wohlfahrt. We thank Achim Kramer for advice and discussion of the manuscript. This work was supported by Helmholtz Health, Metabolic Dysfunction, Helmholtz Alliance ICEMED, DFG KFO218 (all to M.T) and Wellcome Principal Research Fellowship (207462/Z/17/Z), the National Institute for Health and Care Research (NIHR) Cambridge Biomedical Research Centre, the Botnar Fondation, the Bernard Wolfe Health Neuroscience Endowment, the Leducq Foundation grant, and a NIHR Senior Investigator Award (all to I.S.F.).

## Supplementary Figures

**Fig. S1:**
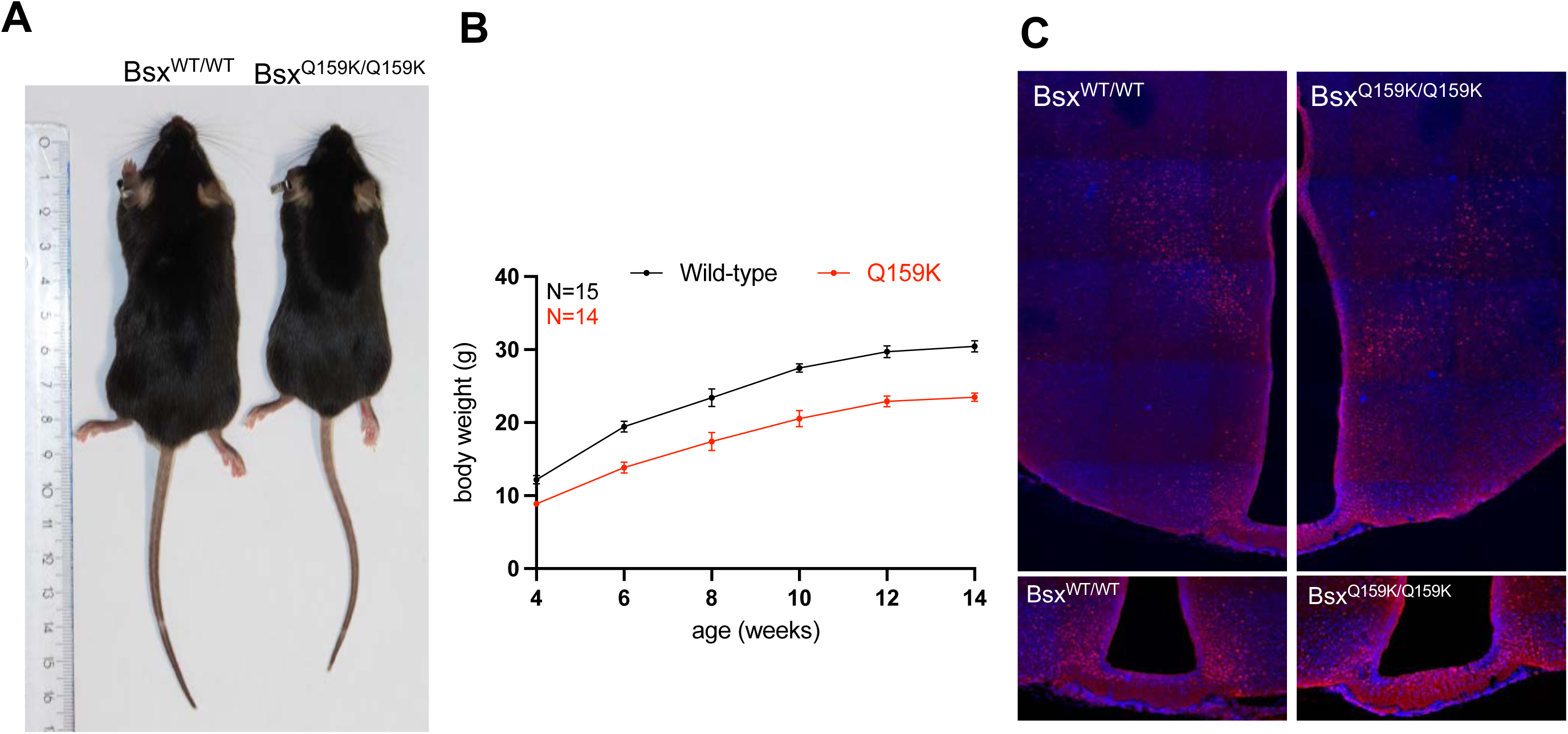
Phenotypic appearance of C57BL/6J Bsx^Q159K/Q159K^ mice. **A)** Representative picture of littermate Bsx^WT/WT^ male and Bsx^Q159K/Q159K^ male mice showing reduction of body size of Bsx^Q159K/Q159K^ compared to their wild-type littermates at 3 months of age. **B)** Growth curve of Bsx^WT/WT^ (N=15) and Bsx^Q159K/Q159K^ (N=14) male mice. Error bars indicate mean and SD. **C)** BSX protein expression is detected by immunohistochemistry in arcuate nucleus and dorsomedial hypothalamus in brains from adult Bsx^WT/WT^ male and Bsx^Q159K/Q159K^ male mice.

**Fig. S2:**
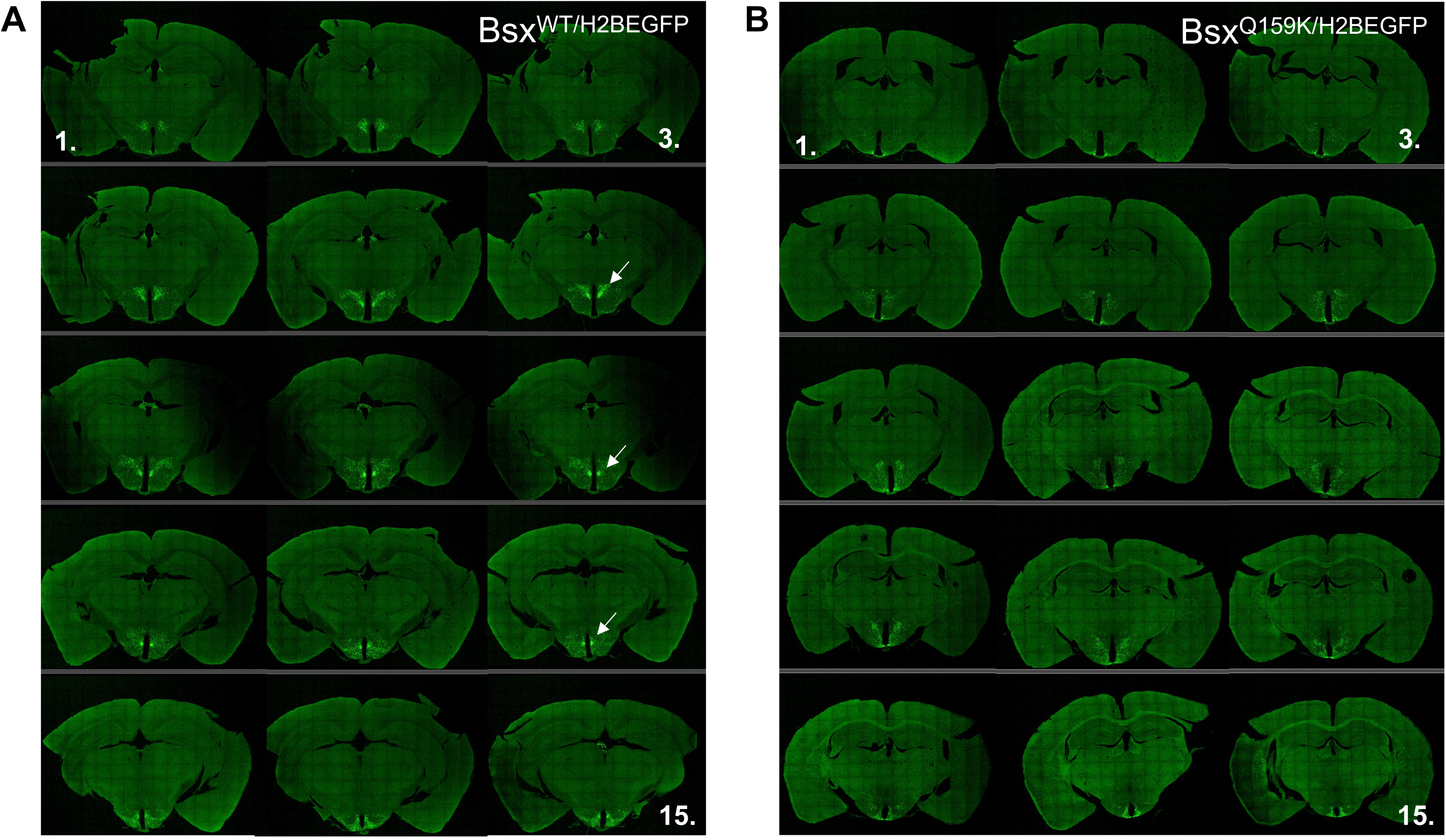
Compact region of the DMH does not form in Bsx^Q159K/H2BEGFP^ mice. **A)** Rostral (1.) to Caudal (15.) adjacent serial 50mm vibratome sections from a Bsx^WT/H2BEGFP^ brain demonstrating compaction of H2BEGFP positive neurons (white arrow) in the dorsomedial hypothalamus (DMH) **B)** Rostral (1.) to Caudal (15.) adjacent serial 50mm vibratome sections from a Bsx^Q159K/H2BEGFP^ brain demonstrating absence of compaction of H2BEGFP expressing neurons in the DMH

**Fig. S3:**
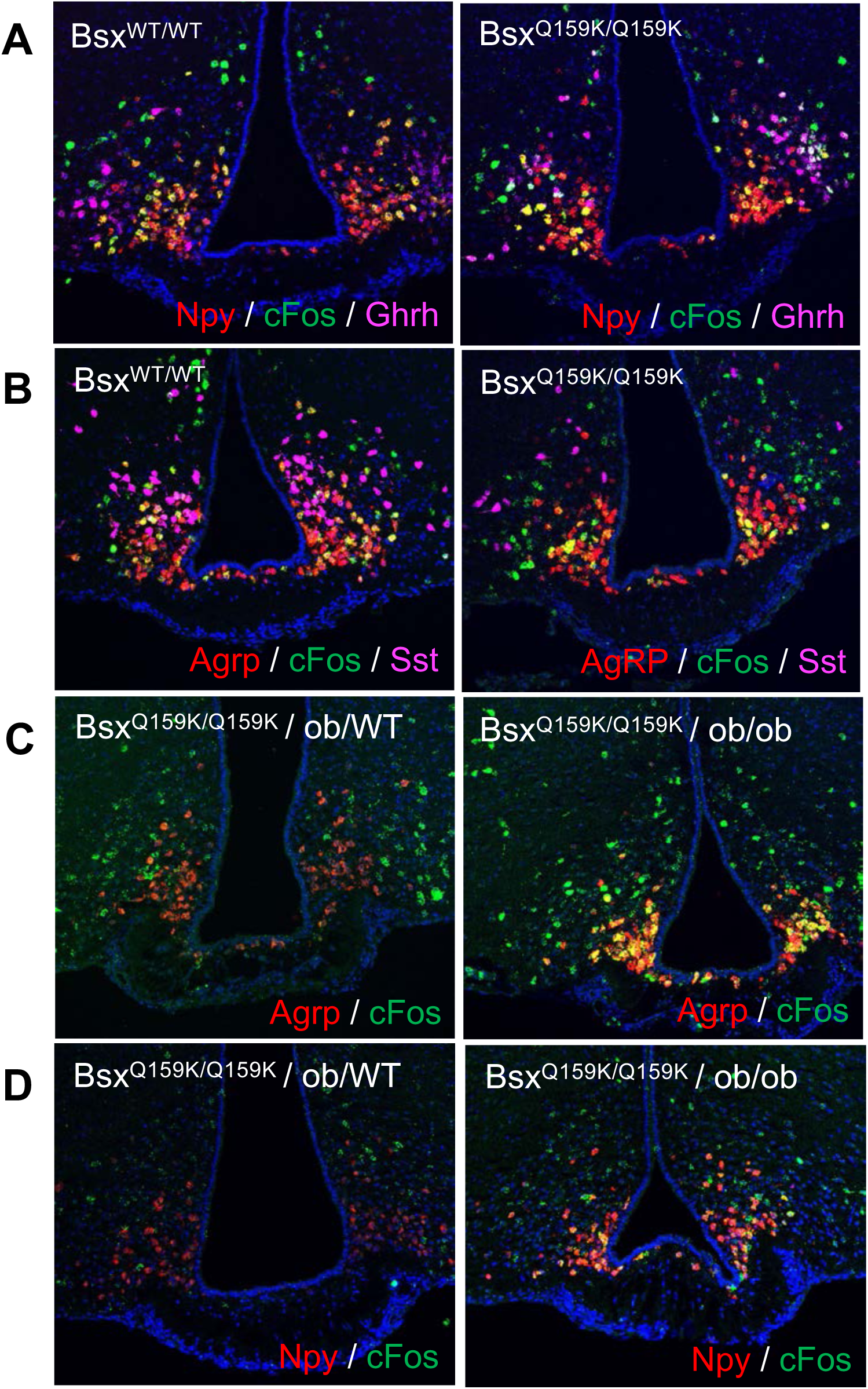
Bsx^Q159K/Q159K^ AgRP/NPY neurons are ghrelin and leptin responsive. **A)** Same amount of c-Fos expression upon ghrelin administration in Bsx^Q159K/Q159K^ and wild-type AgRP/NPY neurons (yellow) with comparable Ghrh neuron numbers in both **B)** Same amount of c-Fos expression upon ghrelin administration in Bsx^Q159K/Q159K^ and wild-type AgRP/NPY neurons (yellow) with less SST neurons (right panel) present in the arcuate nucleus of Bsx^Q159K/Q159K^ mice (independent pair of littermates) **C)** Induction of c-Fos and AgRP expression in Bsx^Q159K/Q159K^ AgRP/NPY neurons (yellow) in the absence of leptin (right panel) **D)** Induction of c-Fos and NPY expression in Bsx^Q159K/Q159K^ AgRP/NPY neurons (yellow) in the absence of leptin (right panel, independent pair of littermates)

**Fig. S4:**
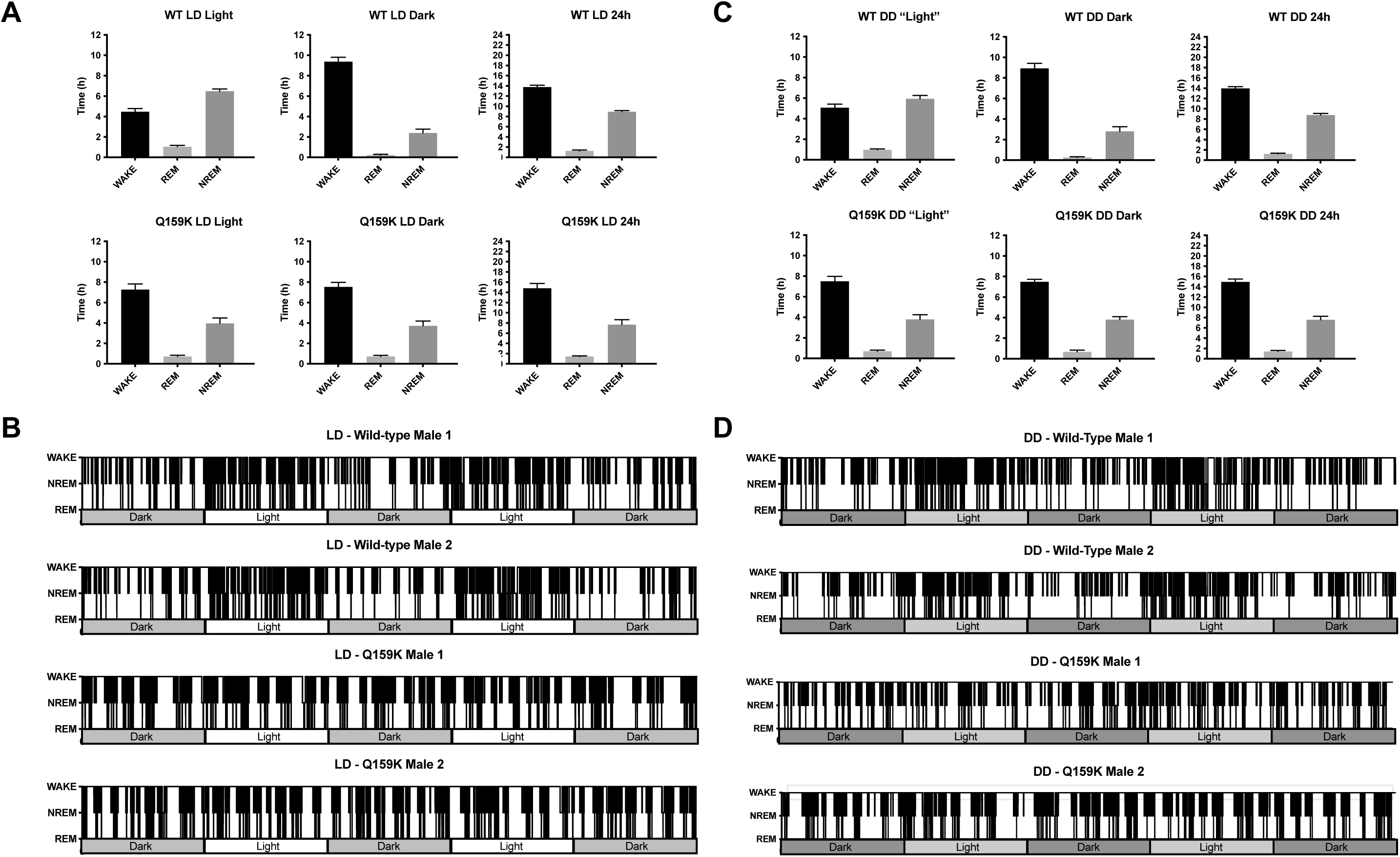
Profound loss of sleep-wake cycles in Bsx^Q159K/Q159K^ mice. A) Wake time, NREM and REM sleep in Bsx^Q159K/Q159K^ mice (N=15) is evenly distributed between light and dark phase in a 12:12 h LD setting with no overall reduction in wake time, NREM and REM sleep compared to wild-type mice (N=11). Error bars indicate SEM. B) 60-h hypnograms of two representative wild-type male mice and two representative Bsx^Q159K/Q159K^ male mice. A clear sleep-wake pattern is obvious in wild-type animals with more NREM and REM sleep and less wake time during the light phase compared to the dark phase. In contrast, Bsx^Q159K/Q159K^ male mice do not show any sleep-wake pattern when comparing light and dark phase or comparing light or dark phase between itself in a 12:12 h LD setting. C) Wake time, NREM and REM sleep in Bsx^Q159K/Q159K^ mice (N=7) is evenly distributed between “light” and dark phase in a constant darkness (DD) setting with no overall reduction in wake time, NREM and REM sleep compared to wild-type mice (N=4). Error bars indicate SEM. D) 60-h hypnograms of two representative wild-type male mice and two representative Bsx^Q159K/Q159K^ male mice. A clear sleep-wake pattern is obvious in wild-type animals with more NREM and REM sleep and less wake time during the “light” phase compared to the dark phase in DD. In contrast, Bsx^Q159K/Q159K^ male mice do not show any sleep-wake pattern when comparing “light” and dark phase or comparing “light” or dark phase between itself in a DD setting.

**Fig. S5:**
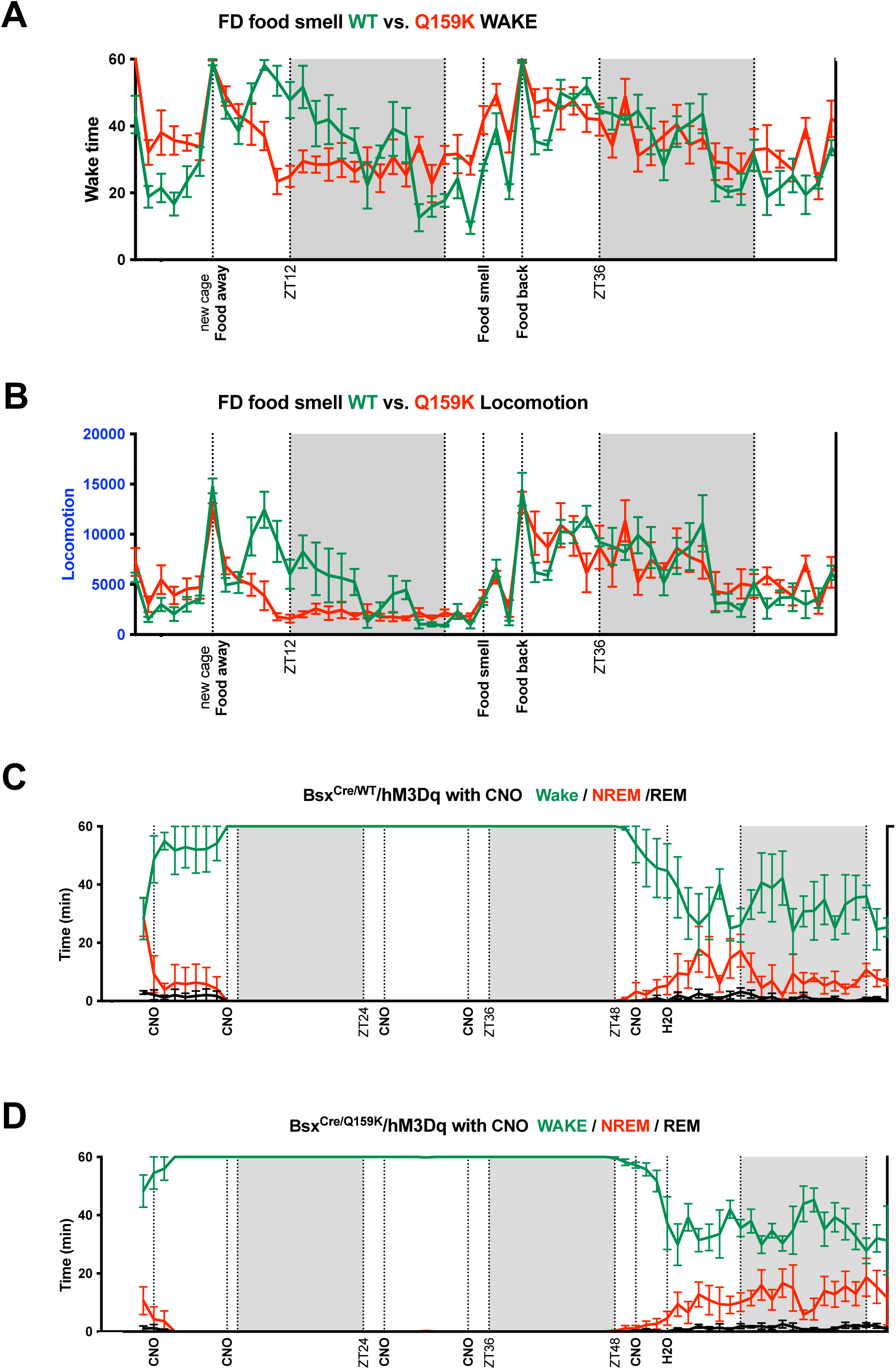
BSX network activity is required and sufficient for food-seeking behaviour. Diagrams are shown as line graphs with mean and error bars (SEM) on the y-axis for wake time (min), NREM (min), REM (min) and locomotion (total beam break counts) respectively and zeitgeber time (ZT) on the x-axis. Grey areas represent time when lights are off. **A)** Cage change and simultaneous food deprivation leads to prolonged increase of wakefulness in Bsx^WT/WT^ male mice (N=5) but only short term increase of wakefulness in Bsx^Q159K/Q159K^ male mice (N=6) whereas food/chow odour exposure after more than 24-h food deprivation elicited short term increase of wakefulness in both Bsx^WT/WT^ and Bsx^Q159K/Q159K^ male mice with wakefulness remaining elevated in both genotypes after feeding. **B)** Locomotor response is impaired under food deprivation in Bsx^Q159K/Q159K^ male mice (N=6) but not Bsx^WT/WT^ male mice (N=5) whereas cage change or food/chow odour exposure elicited short term locomotor activity increase in both genotypes and remained elevated after feeding. **C)** Application of CNO (0.5 mg/ml) through the drinking water to male mice that express hM3Dq in neurons of BSX^WT^ lineage (N=4) induces >36 h of wakefulness with no time spent in NREM or REM stage. **D)** Application of CNO (0.5 mg/ml) through the drinking water to male mice that express hM3Dq in neurons of BSX^Q159K^ lineage (N=5) induces >36 h of wakefulness with no time spent in NREM or REM stage.

**Fig. S6:**
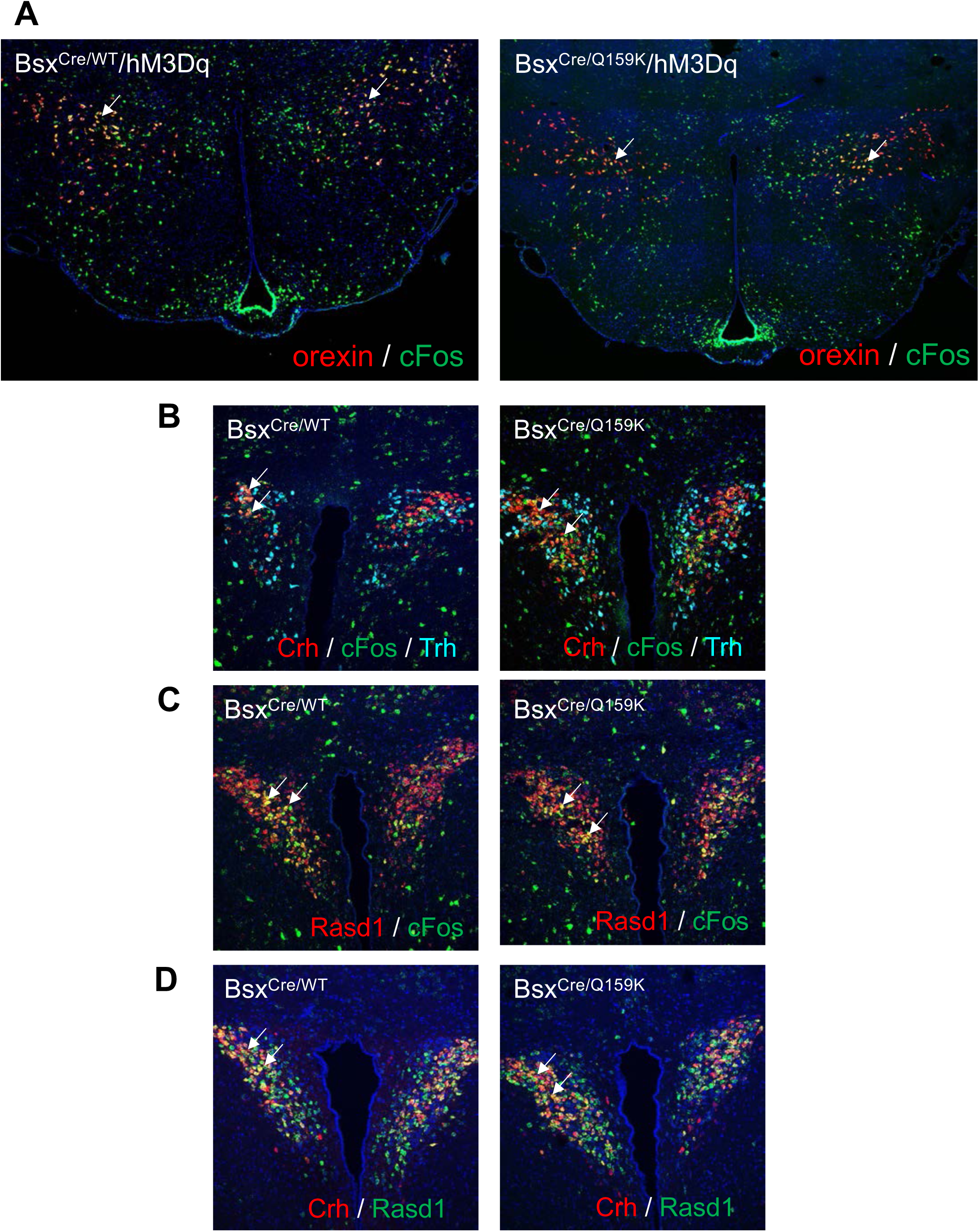
Bsx network activation recruits downstream CRH neurons and orexin neurons. **A)** Representative tile scans of coronal sections of the hypothalamus at the level of the 3^rd^ ventrical after *in situ* hybridization (ISH) with cFos and orexin probes from brains of male mice that express hM3Dq in neurons of either BSX (left) or BSX^Q159K^ (right) lineage 90 minutes after single i.p. injection of 5 mg/kg of CNO. In both cases hypocretin/orexin neurons that do not express Bsx become activated as demonstrated by strong cFos/orexin co-expression in neurons (yellow/white arrows) of the lateral hypothalamus. **B)** Representative *ISH* with Crh, cFos and Trh probes of the PVN from brains of male mice that express hM3Dq in neurons of either BSX (left) or BSX^Q159K^ (right) lineage 90 minutes after single i.p. injection of 5 mg/kg of CNO. In both cases CRH but not TRH neurons that both do not express Bsx become activated as demonstrated by strong cFos/Crh co-expression(yellow/white arrows). **C)** Rasd1 was used as an additional marker to independently demonstrate activation of CRH neurons. Representative *ISH* with cFos and Rasd1 probes of the PVN from brains of male mice that express hM3Dq in neurons of either BSX (left) or BSX^Q159K^ (right) lineage 90 minutes after single i.p. injection of 5 mg/kg of CNO. In both cases PVN neurons that do not express Bsx become activated as demonstrated by strong cFos/Rasd1 co-expression (yellow/white arrows). **D)** Representative *ISH* with Crh and Rasd1 probes of the PVN from brains of male mice that express hM3Dq in neurons of either BSX (left) or BSX^Q159K^ (right) lineage neurons 90 minutes after single i.p. injection of 5 mg/kg of CNO. In both cases CRH neurons that do not express Bsx become activated as demonstrated by strong Rasd1/Crh co-expression (yellow/white arrows).

**Fig. S7:**
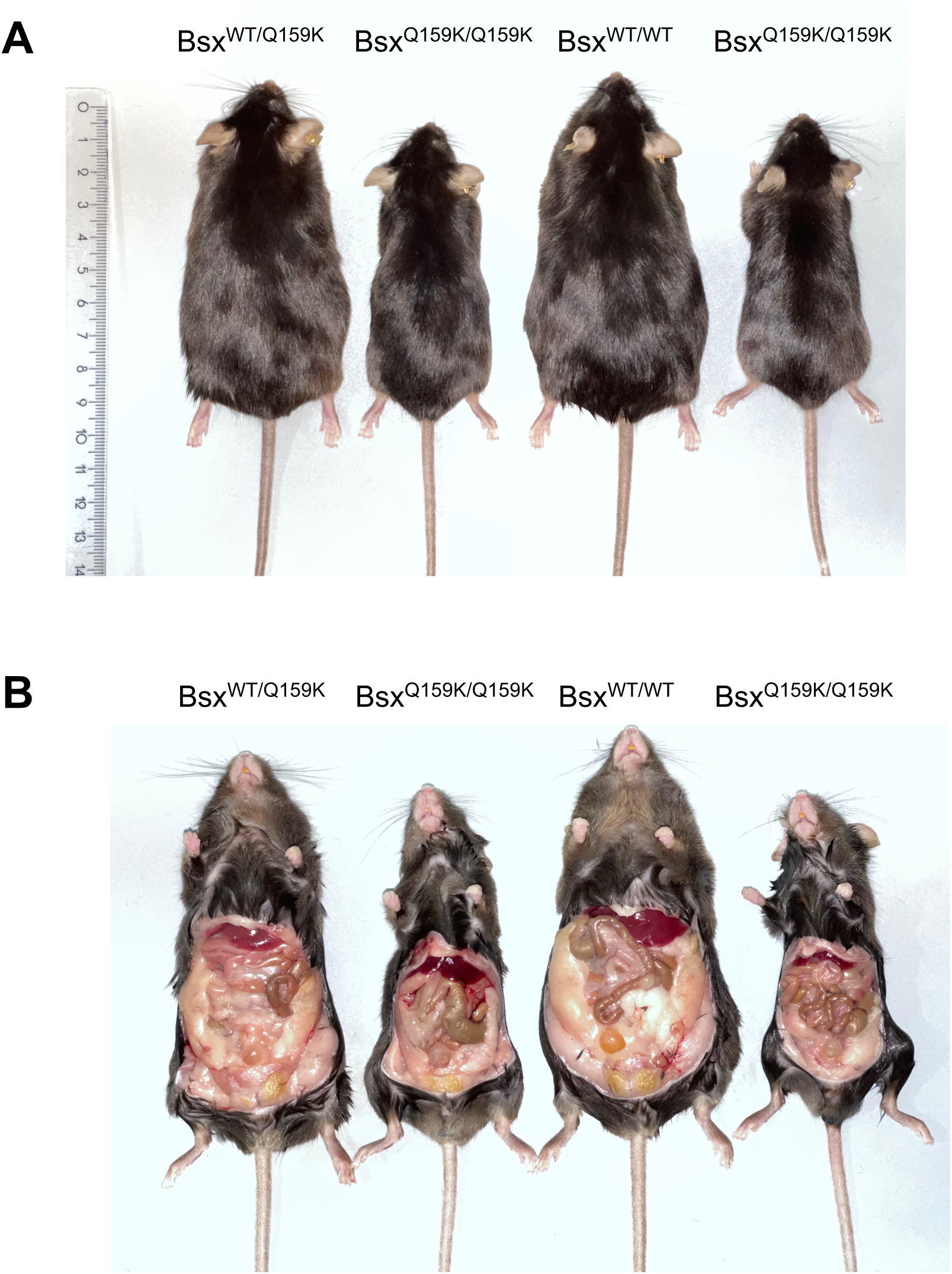
Aged Bsx^Q159K/Q159K^ mice maintain a lean body composition. **A)** Representative picture of 80-week old littermate male mice with all 3 genotypes _Bsx_WT/WT_, Bsx_WT/Q159K _and Bsx_Q159K/Q159K _from a Bsx_WT/Q159K _intercross._ **B)** The same animals with their abdominal wall removed reveals less visceral fat per lean mass in Bsx^Q159K/Q159K^ mice vs. Bsx^WT/WT^ mice visually confirming corresponding NMR measurements.

**Fig. S8:**
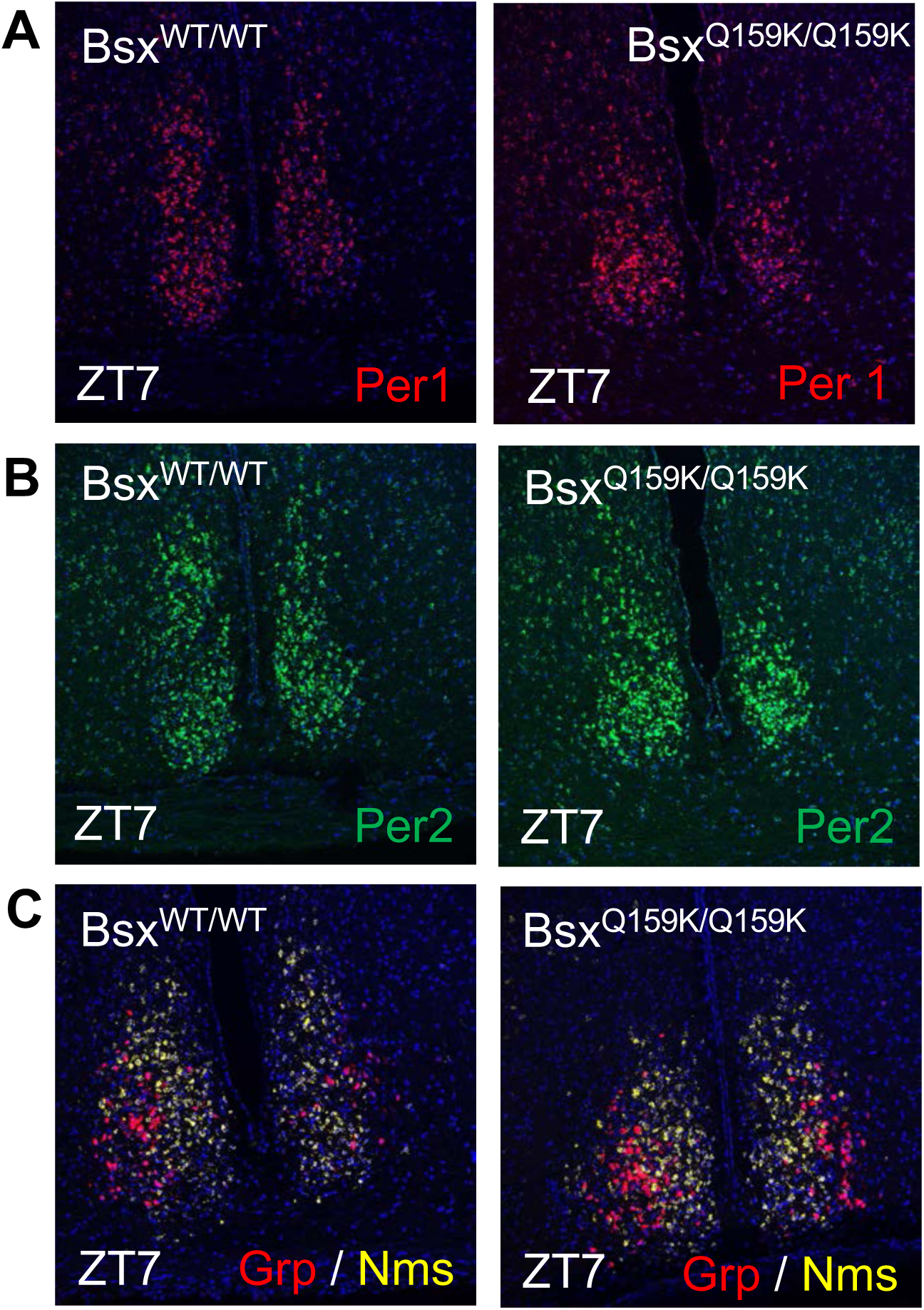
SCN gene expresseion in Bsx^Q159K/Q159K^ mice corresponds to expected ZT time. **A)** Per1 gene expression in the SCN at ZT7 as assed by optical density from in situ hybridization (ISH) is similar in Bsx^WT/WT^ and Bsx^Q159K/Q159K^ during a 12:12 h day night cycle. **B)** Robust Per2 gene expression in the SCN at ZT7 as assed by optical density from in situ hybridization (ISH) is similar in Bsx^WT/WT^ and Bsx^Q159K/Q159K^ during 12:12 h day night cycle. **C)** Grp and Nms gene expression in the SCN at ZT7 as assed by optical density from in situ hybridization (ISH) is similar in Bsx^WT/WT^ and Bsx^Q159K/Q159K^ during 12:12 h day night cycle.

